# Multi-omics dissection of transcriptional and post-transcriptional responses in cyanobacterial high-light adaptation

**DOI:** 10.64898/2026.09.20.752940

**Authors:** Weiyang Chen, Eslam M. Abdel-Salam, Marcel Dann, Caroline Ott, Serena Schwenkert, Dario Leister

**Affiliations:** Plant Molecular Biology, Faculty of Biology, Ludwig-Maximilians-Universität München, Munich, Germany; Computational Systems Biology, RPTU University of Kaiserslautern, Kaiserslautern, Germany; Mass Spectrometry of Biomolecules (MSBioLMU), LMU München, Großhaderner Straße 2-4, 82152 Planegg-Martinsried, Germany; Bio-Inspired Energy Conversion, Technical University of Darmstadt, Darmstadt, Germany

**Keywords:** Cyanobacteria, Synechocystis, High light tolerance, Transcriptomics, Proteomics, NdhF1, EF-G2, Decreased antenna size, Pho regulon

## Abstract

Understanding how photosynthetic organisms acclimate to excess light is essential for the development of robust chassis for synthetic biology approaches aimed at expanding the photosynthetically active absorption spectrum. High-light (HL) tolerant strains were previously generated by laboratory evolution from a laboratory type (LT) strain of *Synechocystis* sp. PCC 6803, with tolerance attributed to a small number of specific point mutations. Key mutations affected the NDH-1L complex F1-subunit (NdhF1_F124L_) and translation elongation factor G2 (EF-G2_R461C_). Reintroduction of these mutations into the LT background was sufficient to confer HL tolerance. However, the mechanisms by which a limited set of point mutations mediates HL tolerance have remained unclear. Here, integrated transcriptomic and proteomic analyses of HL-tolerant strains reveal a coordinated network of responses underlying HL tolerance. The NdhF1_F124L_ mutation increased the accumulation of NDH-1 complex subunits, likely accounting for the previously observed enhancement of cyclic electron flow. In contrast, EF-G2_R461C_ increased the abundance of multiple functional classes of proteins associated with HL tolerance, while post-transcriptionally reducing the level of the phycobilisome linker protein CpcC2, resulting in a decreased antenna size. Integrated analyses further demonstrated that HL tolerance involves transcriptional regulation of protein abundance, including the maintenance of phosphate metabolism. Consistently, overexpression of two genes from the Pho regulon increased HL tolerance. Overall, this study demonstrates how a small number of point mutations in genes with central cellular functions can reprogram the cyanobacterial cell to achieve enhanced tolerance to HL.

## Introduction

Enhancing photosynthetic efficiency is a key objective in efforts to address future challenges in food security, sustainable energy production, and climate change mitigation (Zhu et al., 2010; Slattery and Ort, 2015). One promising strategy is the expansion of the photosynthetically active absorption spectrum through synthetic biology-based approaches and the reconfiguration of the photosynthetic light reactions (Ort et al., 2015; Hitchcock et al., 2022; Leister, 2023). However, excessive solar energy input can lead to photodamage and impaired photosystem function (Powles, 1984). Therefore, the development of robust photosynthetic chassis capable of tolerating increased excitation pressure is required. Such conditions can be experimentally mimicked by exposure to high light (HL) intensities.

Cyanobacteria serve as excellent models for manipulating photosynthesis (Jensen and Leister, 2014; Dann and Leister, 2017). Previous research using adaptive laboratory evolution (ALE) generated *Synechocystis* sp. PCC 6803 (hereafter *Synechocystis*) strains capable of surviving extreme HL conditions (Dann et al., 2021). Two key mutations, phenylalanine (F) 124 to leucine (L) in the F1 subunit of NADH dehydrogenase-like complex (NDH-1) complex (referred to as NdhF1_F124L_) and arginine (R) 461 to cysteine (C) in the translation elongation factor EF-G2 (referred to as EF-G2_R461C_), were found to increase HL tolerance in laboratory-type (LT) background (Dann et al., 2021).

Cyclic electron flow (CEF) plays a vital role in protecting cyanobacteria from reactive oxygen species (ROS) damage during environmental stress, especially under high light, through preventing excess electron accumulation at the PSI acceptor side (Shikanai, 2014). PGR5-dependent pathway acts as the main role for CEF in higher plants, whereas the NDH-1-mediated CEF is the main pathway in cyanobacteria (Ogawa and Mi, 2007; Shikanai, 2014, 2016). In cyanobacteria, the NDH-1 complex is also involved in respiration and CO_2_ uptake, in addition to the CEF, through modulating different subunit compositions (Ogawa and Mi, 2007; Battchikova et al., 2011). NdhD and NdhF subunits have multiple isoforms in cyanobacteria, which are selectively incorporated into different types of NDH-1 complex with the common NDH-1M module (Schluchter et al., 1993; Ohkawa et al., 2000; Zhang et al., 2004; Ogawa and Mi, 2007). NdhF exists in three isoforms, each incorporated into distinct NDH-1 complexes (Schluchter et al., 1993; Ohkawa et al., 2000; Zhang et al., 2004; Ogawa and Mi, 2007). NdhF1, along with NdhD1, is specific to the NDH-1L complex, which is the major type of NDH-1 under normal growth condition and remains stable across different conditions, facilitating CEF around PSI and respiration (Ohkawa et al., 2000; Zhang et al., 2004; Ogawa and Mi, 2007; Battchikova et al., 2011). In contrast, NdhF3 and NdhF4, together with NdhD3 and D4, are components of the NDH-1MS/MS′ complexes, which are mainly involved in CO_2_ uptake, and can also contribute to PSI-CEF and respiratory electron transport through reduction of the plastoquinone pool (Ohkawa et al., 2000; Zhang et al., 2004; Ogawa and Mi, 2007; Battchikova et al., 2011; Bernat et al., 2011; Zhang et al., 2020). Previous studies have demonstrated that HL exposure significantly induces NDH-1 subunit expression, and NDH-1MS is activated during extended HL stress periods (Hihara et al., 2001; Muramatsu and Hihara, 2012; Zhang et al., 2020). Additionally, our prior research revealed that the NdhF1_F124L_ mutation enhances CEF activity, protecting cells against HL stress, shifting the ratio of CEF to CO_2_ uptake activities in favor of CEF, and boosting respiration (Dann et al., 2021).

The translation elongation factor G (EF-G) is responsible for translocating tRNA and mRNA down the ribosome during the translation process. The *Synechocystis* genome contains three genes (*slr1463/fusA*, *sll1098/fusB*, and *sll0830*) that encode different homologues of EF-G (Kojima et al., 2007). EF-Gs have been identified as a primary target within the translational system that becomes inactivated by ROS during photoinhibition (Kojima et al., 2007). Oxidation of EF-Gs inhibits the *de novo* synthesis of the D1 protein, which in turn inhibits the repair of photodamaged photosystem II (PSII) (Kojima et al., 2007; Kojima et al., 2009). The expression of redox-insensitive EF-G1 (encoded by *fusA*) by substituting cysteine (C) 105 with serine has been shown to enhance the repair of PSII through accelerating the synthesis of the D1 protein under strong light (Ejima et al., 2012). EF-G2 (encoded by *fusB*) was significantly targeted during the HL-ALE, leading to the identification of seven independent non-synonymous mutations, and its R461C isoform was validated to enhance the tolerance to HL stress (Dann et al., 2021). It is thus worth investigating whether the addition of a redox-sensitive C in EF-G2_R461C_ can improve its tolerance to oxidative damage under high light conditions, thereby alleviating photoinhibition.

Although it is clear that these two ALE-derived cyanobacterial mutations confer HL tolerance, the underlying mechanisms remain unclear. Therefore, it is important to characterize these HL-tolerant strains to identify the molecular players that can maintain photosynthesis under HL. This study aimed to elucidate the underlying mechanisms of HL tolerance conferred by these mutations through transcriptomic and proteomic analysis of single and double mutant strains. The results provide insights into the functional components involved in HL tolerance and identify common and unique mechanisms across the mutated strains.

## Materials and Methods

### *Synechocystis* strains and growth conditions

Various *Synechocystis* sp. PCC 6803 strains were utilized, including the glucose-tolerant sHL-intolerant laboratory type (referred to LT), the original motile strain (referred to as WT), the HL-adapted monoclonal strain UMMM2 and corresponding strains with point mutations in the LT background as described previously (Dann et al., 2021). To obtain adapted monoclonal strains, single colonies were isolated via streaking on BG11 solid medium from each batch culture following the final round of HL-ALE selection. Subsequently, four clones were selected from each batch as monoclonal strains for whole-genome sequencing and further study. The genome sequence of the LT strain has been deposited at NCBI GenBank under accession numbers CP073017–CP073023 as part of BioProject PRJNA715740, as reported before (Dann et al., 2021). The genomic differences between LT and WT strains have been analyzed in detail in our recent study (Figueroa-Gonzalez et al., 2026), and the corresponding genome sequencing data have been deposited in NCBI GenBank under BioProject PRJNA1228058. All cultures were grown at 23°C in Multi-Cultivator MC 1000-OD devices with an AC-700 cooling unit and warm white LED panel (Photon System Instruments, Brno, Czech Republic). Cells were pre-cultured in liquid BG11 cultures for about a week, then washed and inoculated at OD = 0.05 for photoautotrophic growth under control light (CL, 50 μmol photons per m^-2^ s^-1^), moderate high light (mHL, 700 μmol photons per m^-2^ s^-1^) or strong high light (sHL, 1,200 μmol photons per m^-2^ s^-1^) for one week. For growth assays under varying phosphate conditions, the K HPO concentration was adjusted based on the standard BG11 phosphate concentration of 175 μM. In phosphate-deficient medium, KCl was added at an equivalent potassium concentration to compensate for the potassium omitted by reducing K HPO. Solid media growth used BG11 was supplemented with 0.75% (w/v) bacteriological agar (Carl Roth, Karlsruhe, Germany).

### Generation of *Synechocystis* overexpressors and knock-out mutant

A non-replicative vector lacking the *sacB* selection cassette was employed to generate gene overexpression constructs, as previously described (Dann and Leister, 2019). Whole CDSs of candidate genes were amplified and inserted into the vector behind the *Synechocystis psbA2* promoter using Gibson Assembly® Cloning Kit (New England Biolabs, Ipswich, USA). The resulting plasmids, along with a chloramphenicol resistance (*CmR*) cassette, were inserted into the neutral site *slr0168* of the *Synechocystis* genome. To construct the *cpcC2*-deletion mutant (Δ*cpcC2*), the previously described marker-less gene replacement system was employed (Viola et al., 2014). Approximately 1,000-bp upstream and downstream flanking regions of *cpcC2* were amplified and cloned by Gibson Assembly into a vector derived from the pICH69822 backbone (E. Weber; Icon Genetics GmbH, Halle, Germany), placing them on either side of the *KanR*/*sacB* double-selection cassette. In addition, a second ∼250-bp fragment corresponding to the upstream region of *cpcC2* was inserted between the *KanR*/*sacB* cassette and the downstream flanking region to enable cassette removal during the subsequent negative selection step. The endogenous *cpcC2* locus was replaced with the *KanR/sacB* cassette by homologous recombination upon positive selection with kanamycin.

Transformants were selected on BG11 plates with increasing concentrations of antibiotics (chloramphenicol from 5 to 20 μg/ml and kanamycin from 20 to 100 μg/ml) and confirmed by genomic PCR. The double-selection cassette was removed *via* negative selection on BG11 plate with 5% sucrose.

Both NdhF1_F124L_ and EF-G2_R461C_ were generated by the same marker-less gene replacement system and carry the intended SNP at the corresponding endogenous gene locus (Dann et al., 2021). To confirm complete segregation, the corresponding coding regions were PCR-amplified and verified by Sanger sequencing. The sequencing results confirmed complete segregation of the introduced point mutations. The strains were cryo-preserved immediately after complete segregation, and cultures were regularly refreshed from cryo-stocks to avoid the accumulation of additional mutations.

### RNA isolation and transcriptome sequencing

For RNA extraction, 30 mL of cell cultures (OD_730_ ∼1.0) were harvested after one week of growth. Cells were processed with TRIzol (Invitrogen, Carlsbad, California, USA) at 65°C for 15 min, followed by RNA extraction using phenol-chloroform and isopropanol precipitation (overnight at -20°C). After washing with 75% ethanol, the precipitate was air dried and dissolved in RNase-free water. The RNA was treated with the TURBO DNA-*free*^TM^ kit (Invitrogen, MA, USA) to remove genomic DNA, and then purified using Direct-zol™ RNA MiniPrep Plus columns (Zymo Research, Irvine, California, USA). RNA integrity and quality were assessed using Nano Drop and the Agilent 2100 Bioanalyzer (Agilent, Santa Clara, Calif., USA). Only samples with an RNA Integrity Number (RIN) ≥ 6 were further processed. Ribosomal RNA depletion and RNA-Seq library generation were performed by Novogene Biotech (Beijing, China) using standard Illumina protocols. The RNA-Seq libraries were sequenced on an Illumina HiSeq 2500 system (Illumina, San Diego, Calif. USA) using a 150 bp paired-end sequencing strategy.

### Transcriptome data analysis

RNA-Seq reads were trimmed for adaptor sequences using *Trimmomatic* (Bolger et al., 2014) and sequencing quality was assessed using *FastQC* (http://www.bioinformatics.babraham. ac.uk/projects/fastqc/). Paired-end reads were mapped to the *Synechocystis* sp. PCC 6803 genome (GCF_000009725.1) using HISAT2 (Kim et al., 2015). Read counts were quantified using *featureCounts* (Liao et al., 2014). Principal component analysis (PCA) and UpSet plot were performed in the online web application iDEP (Ge et al., 2018) to determine the reproducibility between three biological replicates and to show group-wise comparisons of the overlapping genes, respectively. One sample (NdhF1_F124L_ at mHL) was widely separated from the other two replicates in the PCA and was omitted from the following analysis. Differentially expressed genes (DEGs) were obtained using *DESeq2* (Love et al., 2014) by applying absolute log_2_ FC > 1 and an adjusted p-value < 0.05 in a contrasting group. FPKM (fragments per kilobase of transcript per million mapped reads) was calculated using the *fpkm* function in *DESeq2*. Genes with FPKM < 1 were determined to be low-expressed genes and removed from downstream analysis.

### Reverse transcription quantitative PCR (RT-qPCR)

The cDNA was synthesized using the iScript cDNA Synthesis Kit (Bio-Rad, Hercules, USA) with 1 µg clean RNA after removing genomic DNA according to the manufacturer’s instructions. The RT-qPCR was conducted on the CFX Connect Real-Time PCR Detection System (Bio-Rad, Hercules, USA) using SsoAdvanced Universal SYBR Green (Bio-Rad, Hercules, USA) and *rnpB* was used as the reference control. All primers used are listed in **Supplementary Data 7**.

### Sample preparation for ^15^N-labelling based quantitative shotgun proteomics

The ^15^N-labelling labeled proteomics was performed as described before (Mühlhaus et al., 2011) with the following modifications. The *Synechocystis* LT strain was grown under constant illumination in BG11 medium containing 9 mM Na^15^NO_3_ as N source and subcultured continuously for 7 days to label the cells with ^15^N. The labeled cells were grown at CL and mHL and then pooled together based on similar cell numbers to generate a reference standard. The ^15^N labelling efficacy of >98% was achieved.

The cell lysis and protein extractions were performed as reported previously (Gandini et al., 2017; Marino et al., 2019). The reference standard from ^15^N-labeled cells was spiked into the unlabeled samples from three biological replicates of each strain under different light conditions at a ^15^N/^14^N ratio of 0.8, based on protein content determined by the BCA protein assay kit (Thermo Fisher Scientific, MA, USA). Proteins were then precipitated in chloroform-methanol mixtures, as previously described (Marino et al., 2019). The pellets were dissolved in 6 M guanidine hydrochloride in 0.1 M HEPES (pH 8.5) containing protease inhibitors (Roche, Mannheim, Germany). The samples were incubated at 60°C for 10 min, sonicated three times for 20 sec, and clarified by centrifugation at 16,000 x g for 15 min. The clarified lysates were transferred to the filter units and digested according to the FASP method as described previously (Wisniewski et al., 2009). The filters were centrifuged at 14,000 x g for 20 minutes and washed with 8 M urea in 0.1 M HEPES buffer (pH 8.5). Bound proteins were reduced by adding 10 mM DTT and incubated at 37°C for 10 min, followed by centrifugation at 14,000 x g for 20 min. Alkylation was performed by adding 0.05 M iodoacetamide for a 5-minute incubation. Samples were centrifuged as before, and filters were washed with 8 M urea in 0.1 M HEPES buffer (pH 8.5) and in HEPES buffer without urea. Proteins were digested with trypsin at 37°C for 16 hours. Digested peptides were purified with home-made C18 stage tips (Rappsilber et al., 2007).

### Liquid chromatography and mass spectrometric analysis

Liquid chromatography-tandem mass spectrometry (LC-MS/MS) was conducted as previously described, with peptides separated over a 40-minute linear gradient of 5–80% (v/v) acetonitrile (ACN) (Marino et al., 2019).

### LC-MS/MS data analysis

Computational analysis of mass spectrometry (MS) measurements was performed using the public Galaxy platform (Afgan et al., 2016). All used tools are available at https://usegalaxy.eu/ under the ‘Proteomics’ and ‘Convert Formats’ sections. The raw files in the proprietary Bruker format generated by the mass spectrometer were converted to the open mzML standard using the msconvert tool (Kessner et al., 2008). Subsequent steps, including ion chromatogram extraction, identification and quantification of labeled (^15^N) and unlabeled (^14^N) peptides, as well as protein-level quantification, were conducted using the ProteomIQon tool chain (v0.0.7).

Peptides were identified from MS/MS spectra by comparing them against the *Synechocystis* reference database employing a target-decoy strategy. Semi-supervised machine learning methods were utilized to integrate multiple search engine scores into a single consensus score (Kall et al., 2007). The results were statistically validated using q-value and PEP-value calculations, with thresholds set at 0.01 for q-values and 0.05 for PEP-values. Peptide abundance was determined by calculating the area under the curve of extracted ion chromatograms. Subsequently, quantified peptide ions were aggregated at the protein level.

Missing values were imputed using the K-Nearest Neighbors (KNN) method within the *imputeLCMD* package at the protein level prior to statistical analysis among replicates. Proteins with high confidence quantitation were filtered by one-way ANOVA with p value <0.05. Then, differentially expressed proteins (DEPs) were determined by applying the fold change >2 in at least in one sample per genotype relative to LT in the three light conditions or in at least one sample in HL relative to CL in the different genotypes.

### Bioinformatic analysis and statistics

Heatmaps with hierarchical clustering were generated using *ComplexHeatmap* v2.10.0 packages (Gu et al., 2016) in R v4.1.0. Functional enrichment analysis was performed in Perseus v2.0.5.0 (Tyanova et al., 2016) with CyanoBase (http://genome.microbedb.jp/cyanobase/) and Gene Ontology (GO) terms. The significance of the enrichment was calculated with Fisher’s exact test. These terms with p value < 0.01 and enrichment fold change >2.0 are considered to be significantly enriched. The enrichment bubble plot was generated using *ggplot2* package in R v4.1.0. The pairwise Pearson’s correlation coefficients between changes of transcript and protein were calculated by running the *ggscatterstats* function in R package *ggstatsplot* (https://github.com/IndrajeetPatil/ggstatsplot).

For fuzzy C-means clustering, the expression profiles of transcripts and proteins were grouped into different clusters using the fuzzy C-means algorithm with the *Mfuzz* package in R v4.1.0. Before clustering, the FPKM of the transcripts and the relative abundance of the proteins were transformed using the *standardize* function in the *Mfuzz* package. The centralized expression pattern in each cluster was represented by a line graph, and each gene or protein was assigned a membership value (0-1) to define its association coefficient with the centralized expression pattern of a given cluster. A cluster membership value >0.9 or 0.6 was used as a threshold for filtering genes or proteins respectively with the best fit to the centralized expression pattern in each cluster.

GraphPad Prism version 10.1.0 software was used for graphs and statistical analyses using one-way ANOVA with post-hoc Tukey HSD test and two-tailed Student’s t-tests.

### Determination of pigment content

After growth, 1 mL *Synechocystis* cells at OD_730 nm_ = 1.0 were centrifuged at 12,000 rpm for 1 min. The pellet was then resuspended in 1 mL 100% methanol and incubated overnight at 4 in the dark. The supernatant containing pigments was centrifuged at 4 for 5 min at 12,000 rpm and analyzed at 470, 665, 720 nm with a spectrophotometer (Ultrospec 2100 pro, Biochrom Ltd., Cambridge, UK). Quantification of chlorophyll and carotenoid content was performed as described before (Zavřel et al., 2015).

### Purification and analysis of phycobilisomes (PBSs)

PBSs were purified as described previously (Glazer, 1988). Briefly, 100 mL cells in exponential phase (OD_730 nm_ ∼1.0) cultured at CL were harvested and washed twice with 0.75 M potassium phosphate (KP) buffer, pH 7.0. Cell pellets were resuspended in 1 ml KP buffer containing protease inhibitor cocktail (Roche, Mannheim, Germany). Cells were disrupted by vortexing with glass beads at 30 Hz for 5 min twice. The broken cell extracts were centrifuged at 4,000 rpm for 20 min, and the supernatants were collected and incubated with Triton X-100 (Carl Roth, Karlsruhe, Germany) at a final concentration of 2% (v/v) for 20 min at room temperature in the darks with gentle shaking. The unbroken cells and debris were removed by centrifugation at 20,000 rpm for 20 min at 15 °C. The dark blue supernatant was loaded onto a 10-40% (w/v) linear sucrose gradient in 0.75 M KP and centrifuged at 30,000 rpm for 16 h at 15 °C. After centrifugation, the blue bands were collected directly with a syringe, and then buffer-exchanged into 200 µL phosphate-buffered saline buffer, pH 7.0, using a 10 kDa Amicon filter. To measure the absorption spectra of purified PBSs, 10 µL PBS solution was mixed with 90 µL phosphate-buffered saline buffer in a 96-well microplate and recorded from 300 nm to 800 nm using a plate reader (Tecan, Maennedorf, Switzerland) at room temperature.

For subunit analysis, 20 µL of PBS solution was mixed with SDS loading buffer and incubated at 42 °C for 2 h. After incubation, the samples were run on an SDS-PAGE gel (12%) with 6M urea and visualized by Coomassie Brilliant Blue staining.

### Protein extraction and immunoblot analysis

*Synechocystis* cells were lysed in a buffer containing 0.4 M sucrose, 10 mM NaCl, 5 mM MgCl_2_ 20 mM Tricine (pH 7.9) and 0.5 mM PMSF with a pre-cooled TissueLyser II homogenizer (Qiagen, Venlo, Netherlands), and the insoluble debris was removed by centrifugation for 10 min at 5,000 g at 4. Protein concentration was measured by BCA assay (Thermo Fisher Scientific, MA, USA). The protein extracts were denatured and loaded onto 10% (w/v) Tris-glycine SDS-PAGE. After gel running, separated proteins were transferred to PVDF membranes (Immobilon-P; Millipore, Burlington, MA, USA) with the Trans-Blot Turbo system (Bio-Rad, Hercules, CA, USA), and visualized by staining with Coomassie Brilliant Blue R-250. The membranes were blocked with 5% (w/v) milk in TBS-T buffer for 2 h at room temperature. After blocking, the membranes were incubated with the primary antibody (anti-HA and anti-AtpB from Agrisera, Vännäs, Sweden) diluted in 3% (w/v) milk with TBS-T overnight at 4.

Protein signals were detected by enhanced chemiluminescence using the SuperSignal™ West Pico PLUS Chemiluminescent Substrate (ThermoFisher Scientific, Waltham, MA, USA) and a Fusion FX ECL reader system (Vilber Lourmat, Collégien, France).

### Detection of intracellular ROS content

For ROS measurements, cells were harvested after cultivation in the Multi-Cultivator, washed, and resuspended in 10 mM phosphate buffer (pH 7.4) to an OD of 0.5. The cell suspensions were incubated with 10 μM 2′,7′-dichlorodihydrofluorescein diacetate (DCFH-DA; Sigma-Aldrich, St. Louis, MO, USA) for 1 h at room temperature in the dark with shaking. Cells were then collected by centrifugation, washed, and resuspended in an equal volume of 10 mM phosphate buffer. DCF fluorescence was measured with a plate reader (Tecan, Maennedorf, Switzerland) using excitation and emission wavelengths of 488 and 530 nm, respectively. For each measurement, the 10 mM phosphate buffer incubated with DCFH-DA was used as blank.

## Results

### HL tolerance of evolved strain UMMM2 and two of its single mutations

In the previous HL-ALE experiment (Dann et al., 2021), the R461C mutation in EF-G2 was first observed after sequential mutagenesis with UV (U) and MMS (M) (UM, **Fig. 1A**). This mutation appeared with high frequency in three of the four monoclonal UMUM strains and in UMMM2 (100% frequency) (**Fig. 1A**) (Dann et al., 2021). In a separate mutagenesis sequence (UMMM), the F124L mutation in NdhF1 appeared at higher frequencies and was detected in two of the resulting monoclonal strains: UMMM2 (0.19% frequency) and UMMM4 (100% frequency) (**Fig. 1A**). Only the UMMM2 strain carries both EF-G2_R461C_ and NdhF1_F124L_ mutations, but with very different allele frequencies (**Fig. 1A**) (Dann et al., 2021). Additionally, the UMMM2 strain contained seven other high-frequency (>80%) mutations, including non-synonymous point mutations in four genes (*rpoC1*, *gltA*, *slr0484*, and *ssr1480*) and frameshifts in three genes (*glcP*, *slr1546*, and *slr1975*).

**Figure 1.**
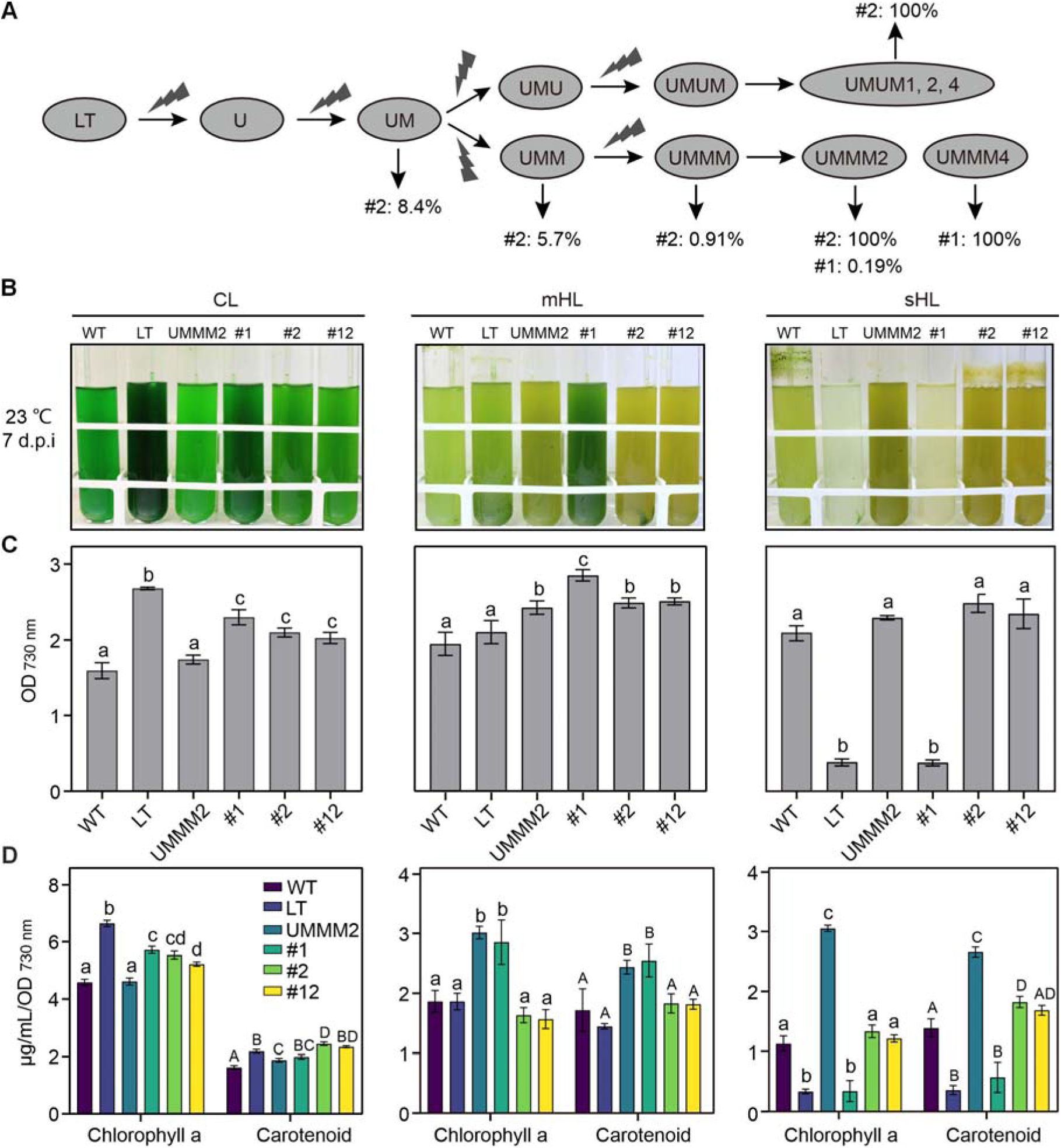
Origin and characteristics of the different strains under different light intensities. **A)** The diagram depicts the mutagenesis process in the HL-ALE experiment, highlighting allele frequencies for two point mutations. U and M represent UV and methyl methanesulfonate (MMS) respectively, which were used as mutagens. Note that the UMMM2 strain harbored seven additional high-frequency (>80%) mutations, including non-synonymous point mutations in four genes (*rpoC1*, *gltA*, *slr0484*, and *ssr1480*) and frameshifts in three genes (*glcP*, *slr1546*, and *slr1975*). **B)** Cultures of various strains (WT, LT, UMMM2, NdhF1_F124L_ (#1), EF-G2_R461C_ (#2), and NdhF1_F124L_+EF-G2_R461C_ (#12)) grown for a week under varying light intensities (CL, mHL, and sHL). Pre-cultures were grown for 7 days at CL at 23°C. The WT strain retains motility, while the LT strain lacks it. All images were captured 7 days post-inoculation (d.p.i.). Notably, UMMM2 is the sole monoclonal strain from HL-ALE containing both NdhF1_F124L_ and EF-G2_R461C_ mutations. **C)** Changes in final optical density (measured at 730 nm). Data points represent mean values with standard deviation (SD), derived from three independent experiments. **D)** Variations in chlorophyll and carotenoid content. The carotenoid values shown here represent total carotenoid content from methanolic extracts. Previous LC-MS/MS analysis identified lycopene, β-carotene, echinenone, zeaxanthin, and myxoxanthophyll-related peaks as the major carotenoid components (Dann et al., 2021). Data points show mean values with SD from three independent experiments. In **C** and **D**, statistical significance (p < 0.05) is indicated by different letters above error bars, as determined by one-way ANOVA with *post-hoc* Tukey HSD test. **ALT TEXT:** Multi-panel figure showing the origin and growth phenotypes of the HL-evolved *Synechocystis* strain UMMM2 and reconstructed site-mutated strains. UMMM2 and reconstructed EF-G2_R461C_-containing strains exhibit improved growth under severe high light, with associated changes in chlorophyll and total carotenoid content.

We compared the HL tolerance of the UMMM2 strain to that of several LT strains containing either the NdhF1 point mutation (NdhF1_F124L_), the EF-G2 point mutation (EF-G2_R461C_), or both (NdhF1_F124L_+EF-G2_R461C_), as well as the LT and the original wild-type motile isolate deposited in the Pasteur Collection (Stanier et al., 1971) (hereafter referred to as WT). Growth assays were performed under three different light conditions: 50 μmol photons m^-2^ s^-1^ (referred to as control light or CL), 700 μmol photons m^-2^ s^-1^ (moderate HL or mHL), and 1,200 μmol photons m^-2^ s^-1^ (strong HL or sHL), at which the LT strain can no longer grow. Our results confirmed the previous findings of Dann et al. (2021) that (i) the NdhF1_F124L_ and EF-G2_R461C_ mutations increase tolerance to HL with a moderate trade-off at CL, (ii) at mHL, NdhF1_F124L_ exhibits superior growth compared to EF-G2_R461C_ and NdhF1_F124L_+EF-G2_R461C_, but is unable to grow at sHL, (iii) at sHL, both EF-G2_R461C_ and NdhF1_F124L_+EF-G2_R461C_ strains are able to grow, and thus the overall behavior of NdhF1_F124L_+EF-G2_R461C_ is similar to that of EF-G2_R461C_, and (iv) the WT strain tolerates both HL conditions, but with a significant trade-off at CL (**Fig. 1B** and **C**, **Supplementary Fig. S1**). Regarding the UMMM2 strain, its pigment content was comparable to NdhF1_F124L_ at mHL and its growth was similar to EF-G2_R461C_ and NdhF1_F124L_+EF-G2_R461C_ at sHL (**Fig. 1B, C** and **D**, **Supplementary Fig. S1**). However, UMMM2 exhibited a higher pigment content than EF-G2_R461C_ and NdhF1_F124L_+EF-G2_R461C_ under HL conditions. At CL, UMMM2 displayed reduced growth and pigment content compared to the three strains with single or combined point mutations (**Fig. 1B, C** and **D**, **Supplementary Fig. S1**). Notably, UMMM2 survived at 2,000 μmol photons m^-2^ s^-1^, whereas EF-G2_R461C_ and NdhF1_F124L_+EF-G2_R461C_ were not viable (Dann et al., 2021).

Taken together, NdhF1_F124L_ and EF-G2_R461C_ are individually sufficient to achieve growth rates under mHL and, in case of EF-G2_R461C_, sHL conditions. However, to tolerate even higher light intensities, the cooperation of the aforementioned additional mutations in UMMM2 seems to be necessary. The inclusion of UMMM2, which was not included in the corresponding analysis in our previous study (Dann et al., 2021), allowed us to directly compare the originally evolved strain with the reconstructed single- and double-point mutants under the same experimental conditions. This comparison revealed that UMMM2 differs from the reconstructed mutant strains in both growth behavior and pigment content. Therefore, the six strains LT, WT, UMMM2, NdhF1_F124L_, EF-G2_R461C_, and NdhF1_F124L_+EF-G2_R461C_ were used as a model system to analyze HL tolerance mechanisms evolved during ALE and their interplay in UMMM2.

### General trends in the transcriptomic HL response

In order to comprehensively identify gene sets associated with HL tolerance, RNA-seq analysis was performed on three biological replicates of WT, LT, UMMM2, NdhF1_F124L_, EF-G2_R461C_, and NdhF1_F124L_+EF-G2_R461C_ grown at CL, mHL and sHL. Transcripts from a total of 3,649 genes were identified in these 54 samples. Principal Component Analysis (PCA) of the global gene expression profile showed that samples at both HL conditions were clearly separated from the profiles at CL, with most strains clustering closely under mHL and sHL (**Supplementary Fig. S2A**), suggesting that some common cellular responses to HL.

Differentially expressed genes (DEGs) were identified (see Materials and Methods), and more DEGs were identified in the three strains with point mutations than in LT, WT and UMMM2 under HL conditions (**Supplementary Fig. S2B**). When comparing the transcriptome profiles of the different strains to that of LT, more DEGs were identified under CL and sHL conditions than under mHL conditions (**Supplementary Fig. S2C**). This suggests that the different genotypes cause substantial differences in the responses to CL and sHL, whereas the responses to mHL appear to be more similar across the different genotypes.

In addition, pairwise overlap analysis showed that WT, UMMM2, and NdhF1_F124L_ generally displayed the highest number of unique DEGs, while EF-G2_R461C_ shared most of its DEGs with NdhF1_F124L_+EF-G2_R461C_ (**Supplementary Fig. S2D-F**). This is consistent with the observation that the double mutation strain is physiologically more similar to EF-G2_R461C_. UMMM2 displayed more unique DEGs than shared ones with NdhF1_F124L_ or EF-G2_R461C_ **(Supplementary Fig. S2D-F**). This suggests that the responses in UMMM2 resulted from the combined effects of its mutations rather than from the sum of individual contributions. Alternatively, it is possible that the impact of NdhF1_F124L_ or EF-G2_R461C_ on the overall transcriptome response of UMMM2 is relatively small. This is supported by UMMM2’s viability at 2,000 μmol photons m^-2^ s^-1^, in contrast to the two point mutation strains.

Next, it was determined whether the accumulation of transcripts for the nine mutated genes in UMMM2, including *ndhF1* and *fusB*, was altered (**Supplementary Fig. S3**). The accumulation of *ndhF1* transcripts was generally decreased at both HL conditions compared to CL in all strains, except for the NdhF1_F124L_ strain, in which down-regulation was already observed at CL (**Supplementary Fig. S3**). The EF-G2_R461C_ mutation did not significantly alter the accumulation of *fusB* transcripts, but reduced *fusB* mRNA accumulation was observed in UMMM2 compared to EF-G2_R461C_ and NdhF1_F124L_+EF-G2_R461C_ (**Supplementary Fig. S3**). Moreover, transcripts for *rpoC1*, *glcP*, and *slr1546* displayed a marked decrease under both HL conditions relative to CL in all strains, except for *rpoC1* in NdhF1_F124L_ (**Supplementary Fig. S3**). Interestingly, the accumulation of transcripts for *slr0484*, *slr1975*, and *ssr1480* showed distinct patterns in UMMM2 compared to LT and three mutants (**Supplementary Fig. S3**).

Taken together, this analysis shows that the response to moderate HL appears to be similar in all genotypes tested, at least at the transcriptome level. The effects caused by NdhF1_F124L_ and/or EF-G2_R461C_ reflect only a portion of the response in UMMM2 due to the presence of seven additional adaptive mutations. In UMMM2, the adaptive mutations not only exchange amino acids in the encoded protein but also affect the expression of other mutated genes at the mRNA steady-state level, indicating that HL adaptation alters expression patterns and regulatory loops.

### Common and genotype-specific pattern in the transcriptomic HL response

A total of 2,366 DEGs were identified in either genotype vs. LT or light conditions vs. CL comparisons and combined for the next analysis (**Supplementary Data Set 1**). Hierarchical clustering analysis grouped the transcript profile of these DEGs into 14 different clusters (**Fig. 2A**), allowing functional enrichment analysis of the genes contained in each cluster (**Fig. 2B**).

**Figure 2.**
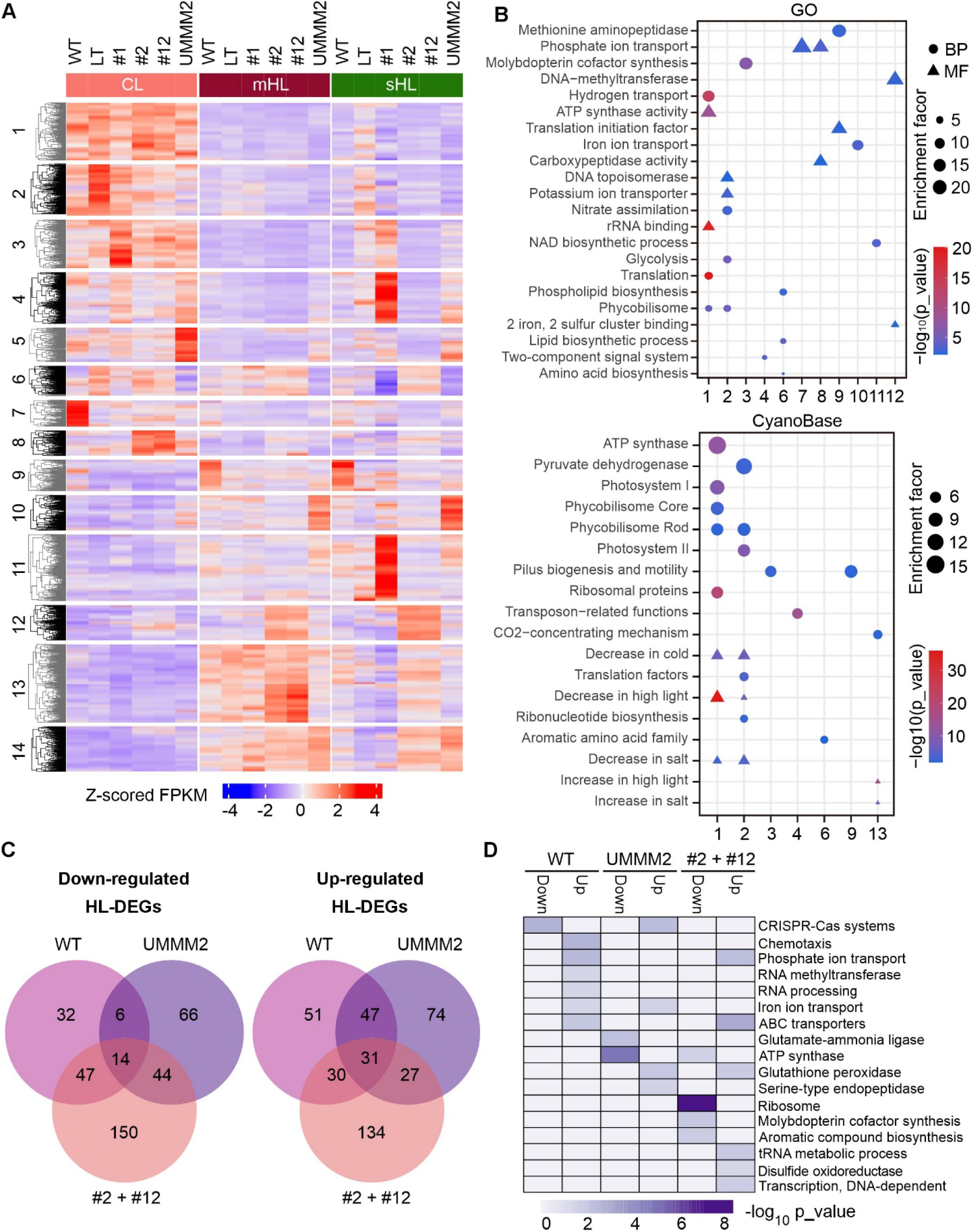
Functional clustering analysis of transcriptome profiles and identification of DEG sets associated with HL tolerance. **A)** Heatmap of DEGs. Fourteen clusters were identified through hierarchical clustering of Z-score normalized FPKM values for 2,366 DEGs, using Euclidean distance and K-means algorithm. **B)** Bubble plot showing non-redundantly enriched functions of different clusters. Functional terms were derived from the CyanoBase database and Gene Ontology (GO). Terms meeting the criteria of enrichment fold change > 2 and p-value < 0.01 (Fisher’s exact test) are displayed. Only clusters with found enrichment are shown. Symbol size indicates enrichment fold change. BP: biological process; MF: molecular function. Terms related to differential expression at HL, cold, and salt stresses (represented by triangles in the lower panel) were obtained from previous publications. **C)** Venn diagrams illustrating the number of shared and unique up- and down-regulated HL-tolerance related genes (HL-DEGs) in HL-tolerant strains: WT, UMMM2, and combined EF-G2_R461C_ (#2) + NdhF1_F124L_+EF-G2_R461C_ (#12). **D)** Heatmap displaying Gene Ontology (GO) terms enriched for up- and down-regulated HL-DEGs in WT, UMMM2, and combined #2 + #12 from **C**. The color scale represents -log_10_ transformed p-values from Fisher’s exact test. **ALT TEXT:** Multi-panel transcriptome analysis showing clustered high-light-responsive genes in LT and HL-tolerant strains. Heatmaps, enrichment bubble plots, and Venn diagrams identify DEG clusters and functional categories associated with high-light tolerance in motile WT, UMMM2, and EF-G2_R461C_-containing strains.

Cluster 1-3 genes were generally down-regulated in both HL conditions, with enrichment in photosynthetic functions (clusters 1 and 2) and translation-related processes (cluster 2) (**Fig. 2A, B**). A decrease in the expression of photosynthesis-related genes is characteristic of long-term or short-term HL stress (Muramatsu and Hihara, 2012) and is thought to alleviate damage caused by excess light. However, the LT and NdhF1_F124L_ strains showed less severe decreases in PSII gene transcripts and notable up-regulation of the three *psbA* genes under sHL (**Fig. 2A, B; Supplementary Fig. S4**). Additionally, ‘Translation’ and ‘rRNA binding’ were enriched in cluster 1 (**Fig. 2A, B**), and transcript accumulation for ribosomal subunits decreased at both HL conditions, except for LT and NdhF1_F124L_ at sHL (**Supplementary Fig. S4**). This suggests that the translation process slowed down when exposed to HL, consistent with the central regulatory role of translation for HL acclimation observed in *Arabidopsis thaliana* (Garcia-Molina et al., 2020).

Genotype-specific transcript patterns were observed in several clusters (**Fig. 2A, B**). For instance, the WT strain displayed unique up-regulation in clusters 7 (enriched for genes involved in ‘Phosphate ion transport’ and ‘Translation initiation factor’) and 9 (enriched for ‘Methionine aminopeptidase’ and ‘Pilus biogenesis and motility’) under CL or the two HL conditions (**Fig. 2A, B**). Similarly, strains with EF-G2_R461C_ mutation showed distinct up-regulation in clusters 8 and 12, enriched for ‘Phosphate ion transport’ and ‘DNA methyltransferase’, respectively (**Fig. 2A, B**). In agreement with our other observations, the general transcript profile of the EF-G2_R461C_ and NdhF1_F124L_+EF-G2_R461C_ strains was highly similar. UMMM2 displayed its own specific transcriptional responses in the clusters 5, 6, and 10, consistent with the low overlap of DEGs between UMMM2 and NdhF1_F124L_ or EF-G2_R461C_. Furthermore, the abundance of gene transcripts in clusters 13 and 14, which are enriched for genes upregulated upon stress conditions, generally increased under both HL conditions in all genotypes except LT and NdhF1_F124L_ at sHL (**Fig. 2A, B**).

These findings demonstrate genotype-specific and light-dependent control of cluster expression, suggesting distinct underlying responses contributing to HL tolerance.

### Gene sets specifically involved in the tolerance to strong HL

Most of the DEGs mentioned above are unlikely to be directly involved in HL tolerance, but may reflect secondary effects. To identify DEGs directly involved in HL tolerance, we exploited the fact that only a subset of our strains was able to grow well under sHL conditions. We categorized strains into sHL-tolerant (WT, UMMM2, EF-G2_R461C_, NdhF1_F124L_+EF-G2_R461C_) and sHL-intolerant (LT, NdhF1_F124L_) groups. By comparing the expression patterns of DEGs, candidate genes directly involved in HL tolerance could be screened depending on whether the expression pattern was unique to the sHL-tolerant group or common to both groups. To identify such expression patterns, we classified all DEGs into distinct clusters based on their expression profiles using the Fuzzy c-means algorithm (Futschik and Carlisle, 2005). We analyzed WT and UMMM2 separately, comparing them to the two intolerant strains (LT and NdhF1_F124L_). Due to their similar responses, EF-G2_R461C_ and NdhF1_F124L_+EF-G2_R461C_ were analyzed together against the intolerant strains.

For WT, we identified 8 distinct clusters (**Supplementary Fig. S5A**). Genes in clusters 7 and 8 were down-regulated in the WT strain in both HL conditions, whereas the two intolerant strains displayed up-regulation in sHL relative to mHL (cluster 7) or similar normalized expression in all three light conditions (cluster 8) (**Supplementary Fig. S5A**). Therefore, genes in clusters 7 and 8 may contain negative regulators of HL tolerance whose down-regulation is relevant for sHL-tolerance in WT. Conversely, genes in cluster 3 and 6 showed up-regulation in the WT strain under mHL conditions compared to CL, and this up-regulation persisted (cluster 3) or even increased (cluster 6) under sHL. In the two intolerant strains, genes were down-regulated at sHL compared to mHL. Therefore, genes in clusters 3 and 6 might contain several positive regulators whose up-regulation is associated with HL tolerance in WT. After filtering with a membership value above 0.9, we identified 99 down-regulated and 159 up-regulated HL-tolerance-related DEGs (HL-DEGs) in WT (**Supplementary Data Set 2**).

Using the same strategy, 8 and 9 different clusters were identified, respectively, after comparing UMMM2 or EF-G2_R461C_/NdhF1_F124L_+EF-G2_R461C_ with LT and NdhF1_F124L_ (**Supplementary Fig. S5B, C**). Using the same routine as described for the WT strain, we again identified clusters containing genes that were either specifically down- or up-regulated with increasing light intensity (**Supplementary Fig. S5B, C**), representing factors that tentatively suppress or promote HL tolerance in UMMM2 or EF-G2_R461C_/NdhF1_F124L_+EF-G2_R461C_. Filtering with a membership value above 0.9 further resulted in 130 down-regulated and 179 up-regulated HL-DEGs in UMMM2, respectively, and for EF-G2_R461C_/NdhF1_F124L_+EF-G2_R461C_ 255 down-regulated and 222 up-regulated HL-DEGs (**Supplementary Data Set 2**).

When comparing the three sets of HL-DEGs sets in the sHL-tolerant strains, it was found that 45% of the down-regulated HL-DEGs and 32% of the up-regulated HL-DEGs from UMMM2 were shared with EF-G2_R461C_/NdhF1_F124L_+EF-G2_R461C_. Furthermore, about 68% of HL-DEGs in WT were shared with UMMM2 or EF-G2_R461C_/NdhF1_F124L_+EF-G2_R461C_, but only a small subset of HL-DEGs were shared by all three (**Fig. 2C**).

To gain insight into the functions of HL-DEGs, we performed their functional annotation based on GO term enrichment. As expected, there was a limited overlap of HL-DEGs among all strains, with no term shared by all three strains together (**Fig. 2D, Supplementary Table S1**). The up-regulated HL-DEGs in WT and EF-G2_R461C_/NdhF1_F124L_+EF-G2_R461C_ showed enrichment in the term ‘Phosphate ion transport’ (**Fig. 2D**), which was found in the cluster analysis within clusters 7 and 8 (**Fig. 2A, B**). Similarly, the term ‘Iron ion transport’ was enriched within up-regulated HL-DEGs in WT and UMMM2, which was enriched within UMMM2-specific cluster 10 (**Fig. 2A, B**). Moreover, enrichment of the term ‘ATP synthase’ was shared by down-regulated HL-DEGs in UMMM2 and EF-G2_R461C_/NdhF1_F124L_+EF-G2_R461C_ (**Fig. 2D**), whereas the terms ‘Ribosome’, and ‘Molybdopterin cofactor synthesis’ were enriched only in down-regulated HL-DEGs of EF-G2_R461C_/NdhF1_F124L_+EF-G2_R461C_ (**Fig. 2D**). Overall, the functional categorization of HL-DEGs from the different sHL-tolerant strains suggests diverse regulatory mechanisms for HL acclimation, likely due to the influence of only a few point mutations.

### Common and genotype-specific control of proteomic responses to HL

To identify proteins associated with the HL tolerance, we characterized the responses at the proteome level using a quantitative shotgun proteomics approach, as previously described (Mühlhaus et al., 2011). Proteins from ^15^N-labeled *Synechocystis* LT cells grown under CL and mHL conditions were spiked into each sample as an internal standard. Data analysis yielded 718 proteins detected in all five samples: WT, LT, NdhF1_F124L_, EF-G2_R461C_, and NdhF1_F124L_+EF-G2_R461C_ samples (**Supplementary Fig. S6A, Supplementary Data Set 3**), excluding the UMMM2 samples due to their high variation. PCA revealed a clear separation between HL and CL conditions, with WT, EF-G2_R461C_, and NdhF1_F124L_+EF-G2_R461C_ clustering closely under both mHL and sHL conditions (**Supplementary Fig. S6B**).

Of the 718 proteins common to all five samples, 347 proteins were identified as differentially expressed proteins (DEPs) (see Materials and Methods, **Supplementary Data Set 4**). The general expression profile of these DEPs was grouped into 12 different clusters using hierarchical clustering analysis. A high similarity between EF-G2_R461C_ and NdhF1_F124L_+EF-G2_R461C_ was also observed at the proteome level (**Fig. 3A**). Some clusters showed expression patterns with clear opposite profiles between the sHL-tolerant and -intolerant strains were evident, such as in clusters 5, 9, 11 and 12 at sHL (**Fig. 3A**), indicating potential candidates of proteins involved in HL tolerance.

**Figure 3.**
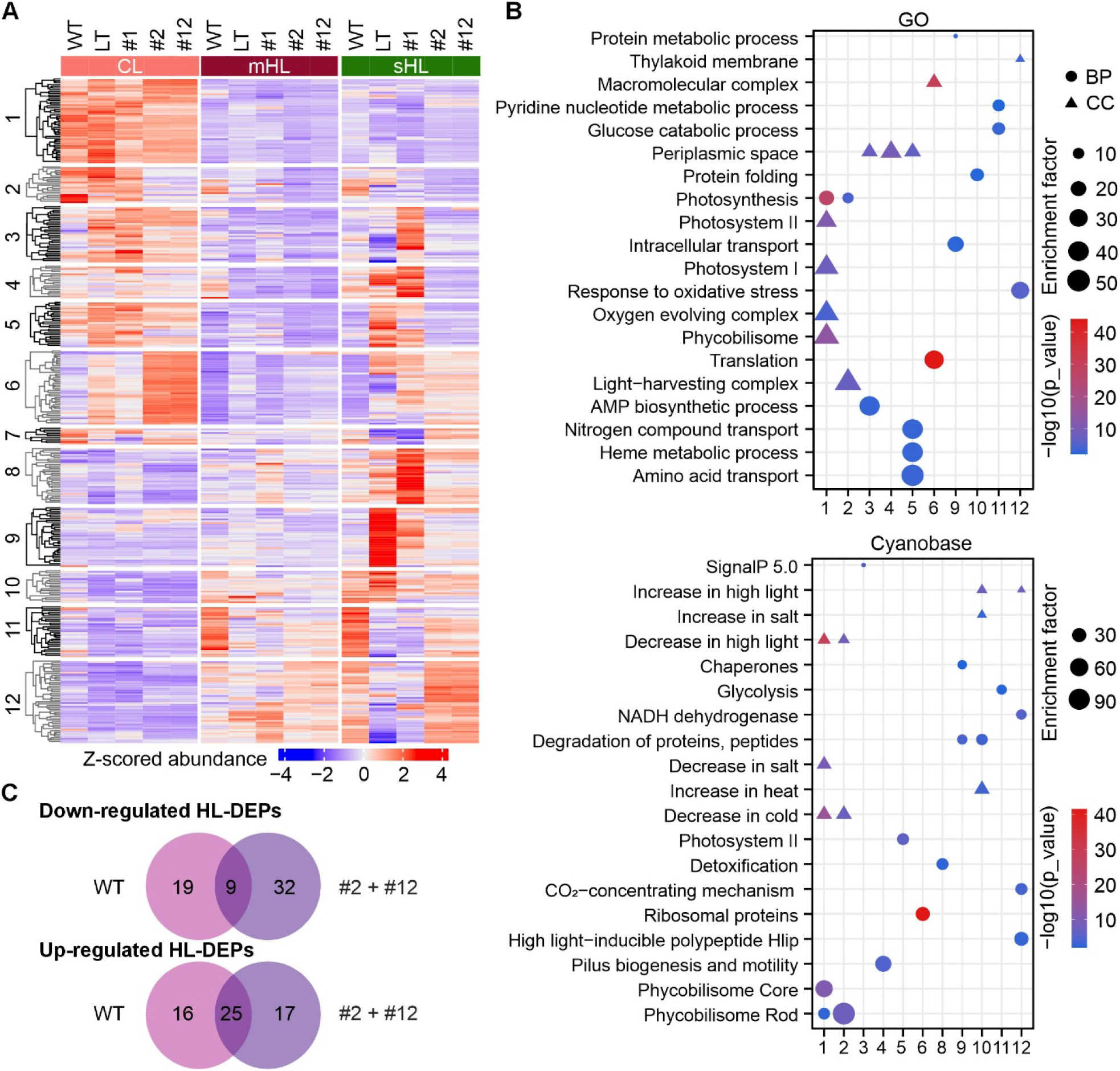
Proteome changes in different strains under varying light intensities and DEP sets associated with HL acclimation. **A)** Heatmap of differentially expressed proteins (DEPs). Twelve distinct clusters were identified through hierarchical clustering of Z-scored relative abundance of 347 DEPs, using Euclidean distance and K-means algorithm. **B)** Bubble plot showing enriched non-redundant functions of different clusters. The methodology used is identical to that described in Fig. 2B. BP: biological process; CC: cellular component. **C)** Venn diagrams illustrating the number of shared and unique up- and down-regulated HL-tolerance related proteins (HL-DEPs) in HL-tolerant strains: WT and combined EF-G2_R461C_ (#2) and NdhF1_F124L_+EF-G2_R461C_ (#12). **ALT TEXT:** Multi-panel proteome analysis showing high-light-responsive protein abundance patterns in LT and HL-tolerant strains. The analysis identifies DEP clusters and enriched functional categories associated with high-light acclimation in motile WT and EF-G2_R461C_-containing strains.

To gain further insight into the functions of DEPs, functional enrichment was performed across these different clusters (**Fig. 3B**). Clusters 1 and 2 consist of proteins that are down-regulated under both HL conditions (**Fig. 3A, B**). Cluster 1 was enriched for the categories ‘Phycobilisome’, ‘PSII’, ‘PSI’, ‘Oxygen evolving complex’, and cluster 2 for ‘Light-harvesting complex’ (**Fig. 3A, B**). These results are consistent with the transcriptional down-regulation of genes associated with these functional categories under HL (see **Fig. 2A, B**). Especially, proteins in cluster 2 are down-regulated in EF-G2_R461C_ and NdhF1_F124L_+EF-G2_R461C_ under CL condition compared to the other three strains, with “Phycobilisome rod” proteins being enriched instead of clustering with the phycobilisome core subunits and other rod subunits in cluster 1 (**Fig. 3A, B**).

Cluster 6 proteins, enriched for ‘Translation’ and ‘Ribosomal proteins’ categories, showed specific up-regulation in the two strains with the EF-G2_R461C_ mutation at CL (**Fig. 3A, B** and **Supplementary S6C**). This closely mirrors the results of our transcriptome analysis (see **Supplementary Fig. S4**). The enhancement of translation may be a coordinated response triggered by the point mutation in EF-G2, as the elongation factor EF-G2 directly participates in the translation process. However, the accumulation of ribosomal proteins was clearly reduced in both strains under HL conditions (**Supplementary Fig. S6C**), again in agreement with the transcriptomic data (see **Supplementary Fig. S4**).

Proteins in clusters 8 and 9 were strongly induced in LT and NdhF1_F124L_ at sHL (**Fig. 3A**) and were enriched in the categories ‘Detoxification’, ‘Degradation of proteins, peptides’, and ‘Chaperones’ (**Fig. 3B**). These functions are designed to assist cells in mitigating the effects of severe stress. This is consistent with the observation that both strains are not able to continue growth at sHL and are likely experiencing protein damage.

Clusters 11 and 12 consist of proteins that are generally up-regulated in both HL conditions in all strains except the sHL-intolerant strains LT and NdhF1_F124L_ (**Fig. 3A**). Metabolic processes and HL stress responses, including ‘High light-inducible polypeptides (Hlips)’, ‘CO_2_-concentrating mechanism (CCM)’, and ‘NADH dehydrogenase’, were enriched (**Fig. 3B**). Cyanobacteria rely on Hlips for survival under HL conditions, and *Synechocystis* has four Hlips (HliA-D) (He et al., 2001). Under HL stress, CO_2_ fixation is enhanced to protect against ROS damage by consuming more reducing power (Muramatsu and Hihara, 2012). Cyanobacteria have evolved sophisticated CCMs to improve the efficiency of carbon fixation, including three bicarbonate transporters: sodium dependent BicA and SbtA, and the high-affinity ATP-binding BCT1 complex, which is encoded by the *cmpABCD* operon (Long et al., 2016). Moreover, the enhancement of NDH-1-mediated CEF plays a key role in preventing the accumulation of excess electrons at the PSI acceptor side, providing more ATP, and thus protecting cyanobacteria against ROS damage under environmental stresses (Battchikova et al., 2011). In our proteomics results, the gradual induction of HliC and HliD, components of the CO_2_-concentrating mechanism (SbtA, CmpA and CcmA), and ten NDH-1 complex subunits were observed in the three sHL-tolerant strains (**Supplementary Fig. S6D, E**). Notably, the NdhF1_F124L_ strain displayed the highest levels of NDH-1 subunits among all strains under mHL but not sHL conditions (**Supplementary Fig. S6E**), as previously observed by immunoblot analysis for NdhK (Dann et al., 2021).

In summary, the protein accumulation pattern provided insight into the putative mechanisms by which the NdhF1_F124L_ and EF-G2_R461C_ mutations enhance HL tolerance. NdhF1_F124L_ mutation promotes increased accumulation of the NDH-1 subunits at mHL, explaining its superior growth (see **Fig. 1**) and higher CEF activity (Dann et al., 2021). The two EF-G2_R461C_ strains are associated with a specific downregulation of phycobilisome rod proteins, a trait previously linked to increased HL tolerance (Kirst et al., 2014; Lea-Smith et al., 2014). Additionally, EF-G2_R461C_ increases ribosomal proteins under CL, and Hlip proteins, CO_2_ concentration proteins and NDH-1 subunits with increasing light intensity. Among the three sHL-tolerant strains, NDH-1 levels were highest at sHL, comparable to the aforementioned NDH levels in the NdhF1_F124L_ strain at mHL, while NDH-1 levels in LT and NdhF1_F124L_ significantly decreased at sHL. This suggests that the level of NDH-1 and, consequently, CEF plays a crucial role in HL-ALE.

### Proteins specifically involved in the tolerance to strong HL

To identify the key proteins involved in HL tolerance, which we refer to as HL-DEPs, we used the fuzzy c-means algorithm to group all DEPs into clusters based on their expression profiles. Then, we identified proteins that might suppress or promote HL tolerance were identified based on their distinctly different expression profiles between sHL-tolerant and -intolerant strains.

After comparing the expression patterns of DEPs from WT and EF-G2_R461C_/NdhF1_F124L_+EF-G2_R461C_ DEPs with the sHL-intolerant strains LT and NdhF1_F124L_, we obtained five and seven distinct clusters, respectively (**Supplementary Fig. S7**). Following the same approach as for the HL-DEGs, DEPs specifically down- and up-regulated in the WT were identified in clusters 5 and 3, respectively (**Supplementary Fig. S7A**). Two clusters (2 and 7) with specific down-regulation with increasing light intensities were identified for EF-G2_R461C_/NdhF1_F124L_+EF-G2_R461C_, and cluster 3 featured DEPs with specific up-regulation in EF-G2_R461C/_NdhF1_F124L_+EF-G2_R461C_ (**Supplementary Fig. S7B**). After applying a membership value threshold above 0.6, we identified 28 down-regulated and 41 up-regulated HL-DEPs in WT. In EF-G2_R461C_/NdhF1_F124L_+EF-G2_R461C_, we detected 41 down-regulated and 42 up-regulated HL-DEPs (**Fig. 3C, Supplementary Data Set 5)**. The PBS rod subunits CpcA, B, C1 and C2 from cluster 2 obtained by hierarchical clustering (see **Fig. 3A**) were identified as down-regulated HL-DEPs, while HliC, SbtA, CmpA, CcmA, and the NDH-1 subunits were found as up-regulated HL-DEPs in EF-G2_R461C_/NdhF1_F124L_+EF-G2_R461C_. Upon comparing the up-regulated and down-regulated HL-DEPs from WT and EF-G2_R461C_/NdhF1_F124L_+EF-G2_R461C_, it was found that nine down-regulated HL-DEPs and twenty-five up-regulated HL-DEPs were shared between both (**Fig. 3C**, **Supplementary Data Set 5**). Only in EF-G2_R461C_/NdhF1_F124L_+EF-G2_R461C_ were PBS rod proteins among the down-regulated HL-DEPs. HliC, SbtA and CcmA were common up-regulated HL-DEPs (**Supplementary Table S2**).

### Correlation of transcriptome and proteome changes

To investigate whether or not transcript and protein changes correlate, we performed a global comparison of the transcriptomic and proteomic datasets. We filtered the datasets using an ANOVA test with a p-value of <0.05, resulting in 660 high-confidence quantified proteins out of 718. As all of these proteins have corresponding transcripts from RNA-seq data, the comparison was performed on these 660 transcript-protein pairs (**Supplementary Data Set 6)**. First, we compared the correlations between the fold changes of transcripts and proteins in each mutant relative to LT at the three different light conditions. The results showed modest positive correlations, with Pearson correlation coefficients ranging from 0.25 for NdhF1_F124L_ vs. LT at sHL to 0.56 for WT vs. LT at mHL. At each light condition, WT vs. LT exhibited the highest coefficient (**Fig. 4A** and **Supplementary Fig. S8A**). This suggests that changes at the protein level can only be partially predicted by changes at the transcript level when comparing different genotypes.

**Figure 4.**
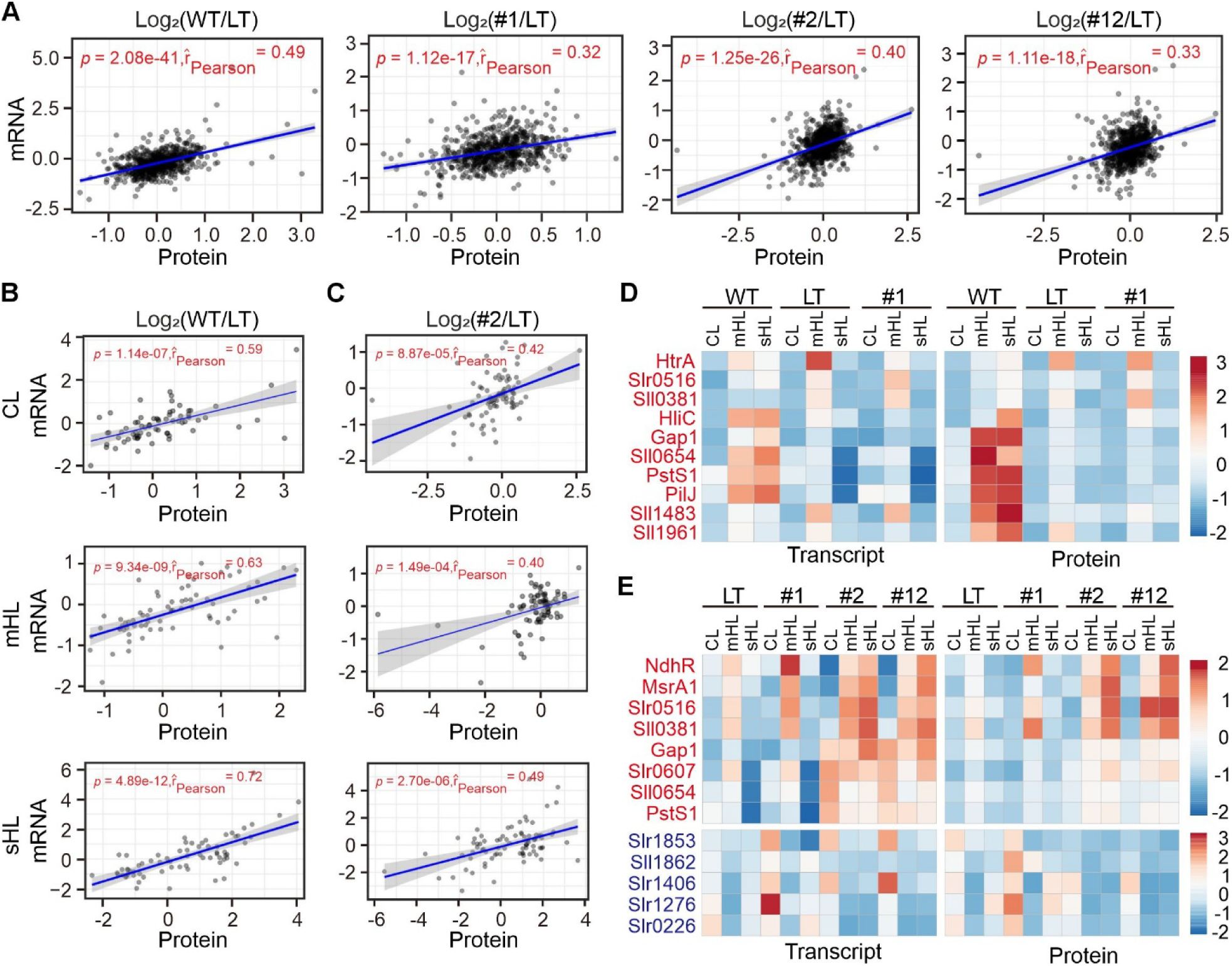
Comparison of transcriptome and proteome responses to different light intensities. **A)** Scatter plots showing correlations between expression changes of all transcript-protein pairs at CL. The x- and y-axes represent log_2_-transformed protein and transcript ratios of WT, NdhF1_F124L_ (#1), EF-G2_R461C_ (#2), and NdhF1_F124L_+EF-G2_R461C_ (#12) relative to LT. Transcript-protein pairs are depicted as gray dots, with best-fit lines in blue. Pearson correlation coefficients (r) and student t-test p-values are provided. **B** and **C)** Scatter plots illustrating correlations between expression changes of HL-DEPs and their corresponding transcripts in WT (**B**) and #2 (**C**). Log_2_-transformed protein and transcript ratios relative to LT under three light conditions (CL, mHL, and sHL) were used for correlation calculations. **D** and **E),** Heatmaps showing abundance changes of transcript-protein pairs shared between HL-DEGs and HL-DEPs in sHL-tolerant WT (**D**) and sHL-tolerant #2 and #12 (**E**) compared to sHL-intolerant strains (LT and #1) under three light conditions (CL, mHL, and sHL). Only up-regulated shared HL-DEGs/DEPs were identified in WT. Genes/proteins from shared up-regulated and down-regulated HL-DEGs/DEPs groups are written in red and blue, respectively. The color bar indicates Z-score normalized FPKM for transcripts and relative abundance for proteins. **ALT TEXT:** Multiple scatter plots showing generally positive but modest correlations between protein and transcript changes in LT and HL-tolerant strains under different light conditions. Heatmaps show that only a small subset of high-light-responsive genes and proteins overlap in WT and EF-G2_R461C_-containing strains.

Higher coefficients were observed when comparing the correlations between fold changes of transcript-protein pairs from CL to mHL or sHL in the same genotype (**Supplementary Fig. S8B**), suggesting that proteome responses to HL are more predictable from transcriptome data when analyzing the effects of different light conditions in the same strain. An exception was found in the very low correlation (0.20) obtained for LT in the sHL vs. CL comparison (**Supplementary Fig. S8B**), possibly due to the cellular stress and its growth arrest at sHL.

Furthermore, when limiting the analysis to the HL-DEPs in WT and EF-G2_R461C_, the correlation coefficients for WT vs. LT and EF-G2_R461C_ vs. LT, respectively, were found to be higher for the three light conditions than for the overall comparisons that included all DEPs (**Fig. 4B, C**). Notably, only for a limited proportion of the HL-DEPs was the corresponding DEG also identified as HL-DEG. For WT, none of the DEGs corresponding to the 28 down-regulated HL-DEPs were identified as down-regulated HL-DEGs, and only 10 of the 41 up-regulated HL-DEPs were also identified as up-regulated HL-DEGs (**Fig. 4D**). Similarly, for EF-G2_R461C_, the proportion of HL-DEPs with a DEG counterpart that was also a HL-DEG was rather small. Only 8 out of 43 up-regulated HL-DEPs and 5 out of 41 down-regulated HL-DEPs were identified as HL-DEGs, respectively (**Fig. 4E**). These results suggest that only a portion of the HL response involves transcriptional regulation of protein accumulation.

### Transcript-level independent regulation of CpcC2 accumulation may play a role in HL tolerance

The limited correlation between transcript and protein changes, along with the low overlap between HL-DEGs and HL-DEPs, suggests that factors beyond steady-state mRNA levels may influence HL-tolerance. Given that the EF-G2_R461C_ mutation affects an elongation factor, some protein accumulation changes in this strain might be directly caused by the mutation itself. To identify such cases, we searched for clear discrepancies between transcript and protein changes, focusing on anticorrelations. By plotting log_2_ (EF-G2_R461C_ vs. LT) values for transcript and protein changes across three light conditions, we identified 25 instances where proteins were at least 3-fold up- or down-regulated while corresponding transcripts were either oppositely or minimally regulated. Of these, 14 were HL-DEPs in EF-G2_R461C_ (**Supplementary Data Set 6)**.

To confirm whether these transcript-level-independent regulations were related to HL-tolerance in EF-G2_R461C_, we investigated one candidate, CpcC2, a down-regulated HL-DEP in the EF-G2_R461C_ and NdhF1_F124L_+EF-G2_R461C_ strains. In EF-G2_R461C,_ compared to LT, much less CpcC2 protein was found with only a slight decrease under CL and mHL light conditions and a moderate decrease under sHL in transcript levels **(Fig. 5A)**. CpcC2 is a phycobilisome (PBS) protein, part of the light-harvesting complex in *Synechocystis*. The *Synechocystis* PBS consists of a core allophycocyanin (APC) region and six radiating rods, each composed of three stacked disc-shaped phycocyanin (PC) hexamers (Arteni et al., 2009). Linker proteins connect the discs; the disc proximal to the APC core is connected *via* CpcG1 or CpcG2, the middle disc *via* CpcC1, and the distal disc *via* CpcC2 (Arteni et al., 2009). The deletion of CpcC2 and CpcC1 results in the loss of the distal disc and middle disc on the rod and a smaller antenna size. However, it has no significant effect on photoinhibition or growth at higher light intensities (Lea-Smith et al., 2014). A reduction in absorbance at 625 nm specific to PC was observed in EF-G2_R461C_ and NdhF1_F124L_+EF-G2_R461C_ in whole-cell absorption spectra (Dann et al., 2021). To confirm this, the PBSs were purified and resolved by sucrose density gradient. The results showed that PBSs from EF-G2_R461C_ and NdhF1_F124L_+EF-G2_R461C_ shifted at higher position (**Fig. 5B**), indicating a smaller size. The PBSs from UMMM2 displayed a higher shift than LT and NdhF1_F124L_, but somewhat lower than in EF-G2_R461C_ and NdhF1_F124L_+EF-G2_R461C_ (**Fig. 5B**). This suggests that the R461C mutation in EF-G2 in UMMM2 also resulted in its reduced PBSs. A similar trend was also detected from the absorption spectrum of these purified PBSs (**Fig. 5C**). The subunit profiles of these purified PBSs were also analyzed. The results indicated that the CpcC2 protein in EF-G2_R461C_ and NdhF1_F124L_+EF-G2_R461C_ was almost undetectable (**Fig. 5D**), which is consistent with the proteomics result. A significant decrease of CpcC2 was also detected in UMMM2 PBSs, but not as drastic as in EF-G2_R461C_ and NdhF1_F124L_+EF-G2_R461C_ (**Fig. 5D**). In addition, CpcC1 showed a similar but less pronounced down-regulation compared with CpcC2 (**Fig. 5A and D**). Taken together, it appears that the EF-G2_R461C_ mutation leads to a reduced antenna size through decreased accumulation of the CpcC2 linker protein, independently of changes in steady-state transcript levels.This mechanism may contribute to the enhanced HL-tolerance observed in these strains.

**Figure 5.**
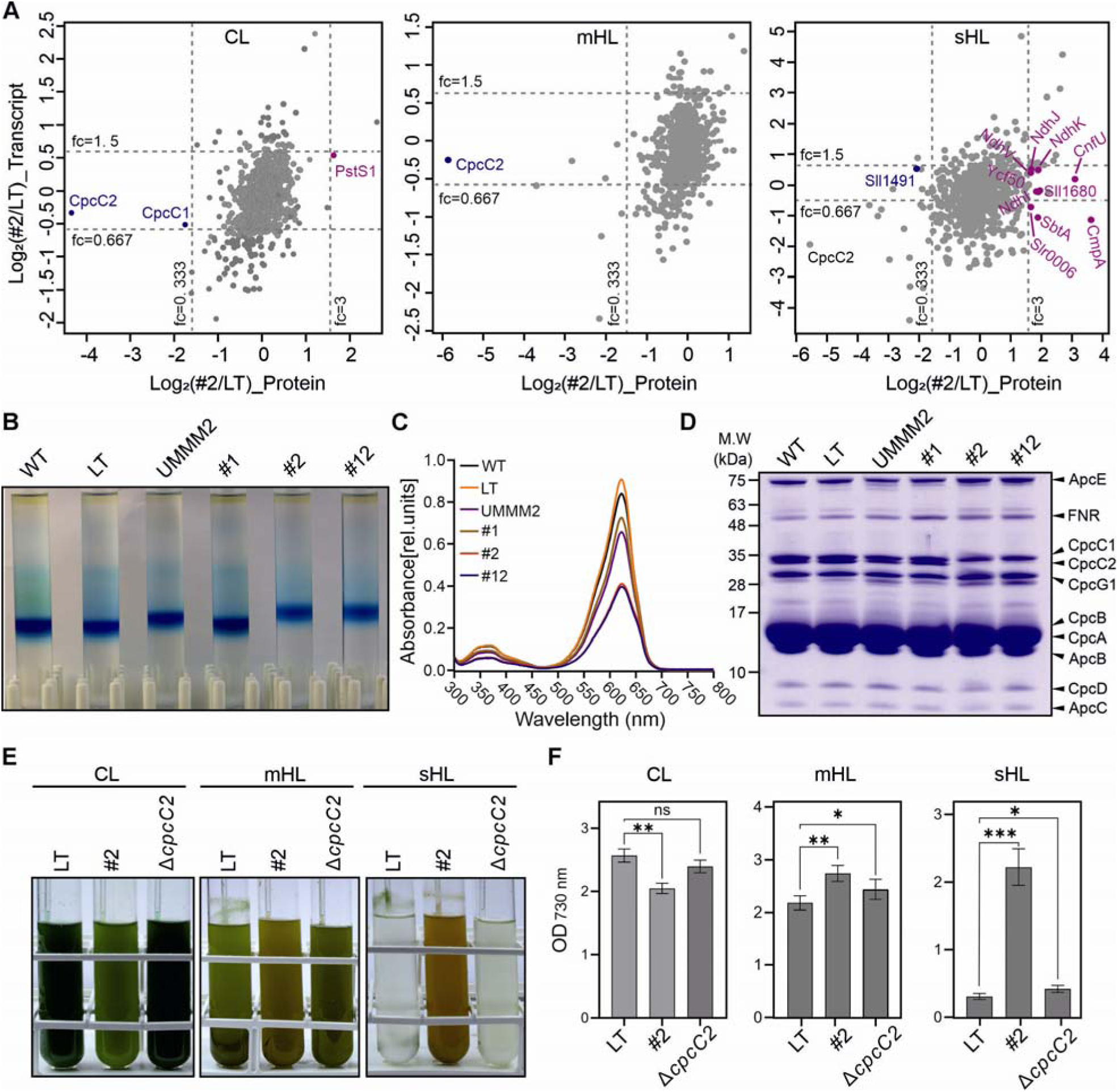
Transcript-independent regulation of CpcC2 contributes to HL tolerance in EF-G2_R461C_ strains. **A)** Scatter plot comparing log_2_-transformed ratios of transcripts and proteins in EF-G2_R461C_ (#2) relative to LT (#2/LT) under three light conditions (CL, mHL, sHL). Highlighted are proteins with markedly different changes from their corresponding transcripts (at least 3-fold up- or down-regulated at protein level, while either oppositely or no more than 1.5-fold regulated at transcript level) across all light conditions. Among these, up-regulated and down-regulated HL-DEPs in #2 are labeled in pink and blue, respectively. **B)** PBS size analysis using sucrose gradient. The blue band indicates purified PBS positions, with higher positions suggesting decreased PBS size. Strains: WT, LT, UMMM2, #1 (NdhF1_F124L_), #2 (EF-G2_R461C_), #12 (NdhF1_F124L_+EF-G2_R461C_). Image shown is representative of three independent experiments. **C)** Absorption spectrum of purified PBSs at room temperature. Peak absorbance at 625 nm corresponds to phycocyanin (PC). Spectra normalized at 300 nm. Data represents means from three independent experiments. **D)** Protein composition of purified PBSs. SDS-PAGE separation of proteins from purified PBSs, stained with Coomassie Brilliant Blue. Gel shown is representative of three independent experiments as in **B**. **E)** Culture images of LT, EF-G2_R461C_ (#2), and Δ*cpcC2* grown for 7 days under various light intensities (CL, mHL, and sHL). Images shown are representative of three independent experiments. **F)** Optical density (OD_730 nm_) of LT, EF-G2_R461C_ (#2), and Δ*cpcC2* cultures after 7 days of growth. Data shows mean ± SD from three independent experiments as in **E**. Statistical significance was determined using two-tailed Student’s t-test. *: p < 0.05, **: p < 0.01, ***: p < 0.001. **ALT TEXT:** Multiple scatter plots in panel A illustrate the discrepancy between transcript and protein abundance of CpcC2 in EF-G2_R461C_ relative to LT under three light conditions. Sucrose-gradient, absorption-spectrum, and protein gel analyses indicate reduced phycobilisome antenna size and CpcC2 abundance in EF-G2_R461C_, while growth assays of Δ*cpcC2* show a modest improvement under high-light conditions.

To further test this speculation, we generated a *cpcC2* deletion mutant, Δ*cpcC2*, in the LT background using the previously established markerless gene replacement system (Viola et al., 2014). In this mutant, only the *cpcC2* locus was removed, without disrupting other genes in the same *cpc* operon (**Supplementary Fig. S9A**). Growth analyses showed that Δ*cpcC2* achieved higher final cell densities and pigment contents than LT after 7 days of cultivation under mHL and sHL, without an apparent growth trade-off under CL (**Fig. 5E**, **F**; **Supplementary Fig. S9B, C**). However, its HL tolerance was still substantially lower than that of EF-G2_R461C_, particularly under sHL conditions. Therefore, these results suggest that the loss of CpcC2 contributes to improved HL acclimation, but is not sufficient to fully recapitulate the strong sHL-tolerant phenotype of EF-G2_R461C_.

### Overexpression of members of the phosphate regulon improves HL tolerance

From the small set of overlapping HL-DEGs and HL-DEPs that show correlation of transcript and protein levels (see **Fig. 4D, E**), 4 up-regulated ones (Sll0381, Gap1, PstS1, and Sll0654) were shared between WT and EF-G2_R461C_. PstS1 is a member of the ABC-type inorganic phosphate (P_i_) transport system in cyanobacteria, and its expression significantly increases when cells are challenged with P_i_ limitation stress (Pitt et al., 2010). PstS1 and PstS2 bind P_i_ and are encoded by two separate *pst* gene clusters, along with other P_i_ transport components (Pitt et al., 2010). Interestingly, all *pst* genes showed increased transcript accumulation in both WT and EF-G2_R461C_ (**Supplementary Fig. S10A**), and the term ‘Phosphate ion transport’ was enriched among up-regulated HL-DEGs in both WT and EF-G2_R461C_ (see **Fig. 2D**). Another protein in this set of four is Sll0654, which is a putative alkaline phosphatase. Sll0654, along with these P_i_ transporters, belongs to the phosphate (Pho) regulon, which is related to P_i_ metabolism and is strongly induced by P_i_ limitation (Suzuki et al., 2004; Pitt et al., 2010). Interestingly, the Pho regulon is up-regulated when shifted to HL (Bhaya et al., 2000; Suzuki et al., 2004; Pitt et al., 2010). In both the LT and NdhF1_F124L_ strains, the expression of the Pho regulon was lower compared to the sHL-tolerant strains and was down-regulated under sHL conditions (**Supplementary Fig. S10A**), suggesting that tolerance to sHL might be linked to the level of expression of the phosphate regulon.

Both Sll0654 and PstS1 showed increased or maintained expression at both transcript and protein levels in WT and the two strains carrying the EF-G2_R461C_ mutation with increasing light intensities, in contrast to the two sHL-intolerant strains. A direct way to test their effect on HL tolerance is to overexpress them. Therefore, we generated overexpression (OE) lines PstS1-OE and Sll0654-OE and validated the up-regulation of both (**Supplementary Fig. S10B-D**). Growth comparisons with LT under CL, mHL, and mHL’ (500 μmol photons m-2 s-1) showed that LT initially grew faster (**Fig. 6B**). However, at mHL and mHL’, LT growth retarded after 48h, while PstS1-OE and Sll0654-OE outperformed LT, in terms of final cell concentration and pigment content after 7 days of cultivation (**Fig. 6A-C, Supplementary Fig. S10E**). However, neither of the OE lines continued to grow at sHL, similar to LT (**Fig. 6A-C, Supplementary Fig. S10E**). Therefore, these results suggest that an increase in P_i_ transport and metabolism would be a conserved and universal regulatory response for enhancing HL tolerance, albeit with a growth trade-off at low light conditions, as already observed in WT, UMMM2, and EF-G2_R461C_ cells.

**Figure 6.**
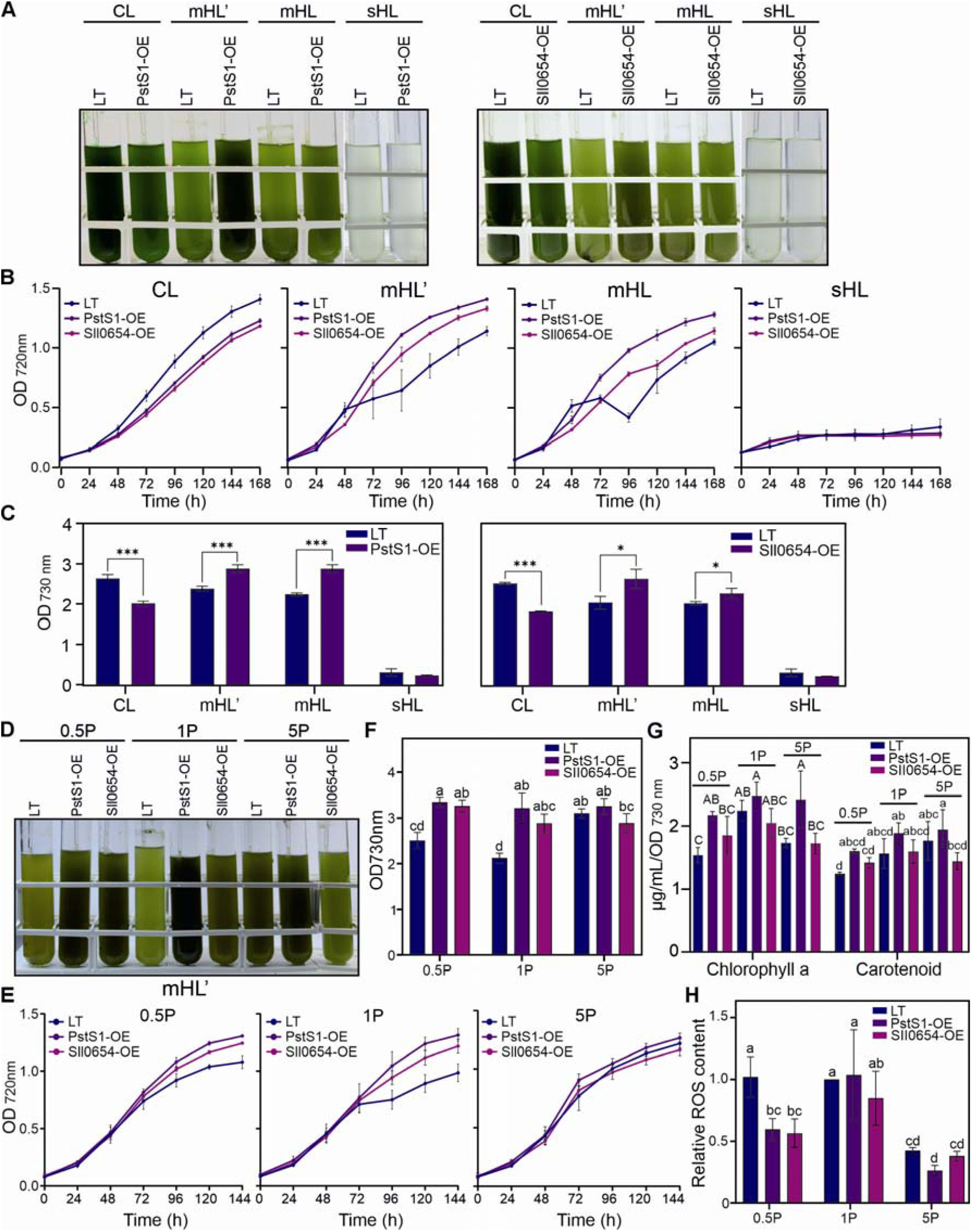
Overexpression of PstS1 and Sll0654 enhances high light tolerance. **A)** Culture images of LT, PstS1-OE, and Sll0654-OE grown for 7 days under various light intensities (CL, mHL’, mHL, and sHL). mHL’ represents 500 μmol photons m^-2^ s^-1^. Pre-cultures were grown for 7 days at CL at 23°C. Images shown are representative of three independent experiments. **B)** Growth curves of LT, PstS1-OE, and Sll0654-OE under CL, mHL’, mHL, and sHL conditions (mHL’: 500 μmol photons m^-2^ s^-1^). Optical density (OD_720 nm_) was measured hourly using the built-in real-time monitor system in Multi-Cultivator OD-1000 devices. For clarity, values at 24-hour intervals were plotted. Data shows mean ± SD from three independent experiments as in **A**. **C)** Optical density (OD_730 nm_) of LT, PstS1-OE, and Sll0654-OE cultures after 7 days of growth. Data shows mean ± SD from three independent experiments as in **A**. Statistical significance was determined using two-tailed Student’s t-test. *: p < 0.05, **: p < 0.01, ***: p < 0.001. **D -G),** Growth summary of LT, PstS1-OE, and Sll0654-OE grown for 6 days under mHL’ with various phosphate (P_i)_ intensities. 1P represents standard BG11 medium containing 175 μM K HPO ; 0.5P represents low-phosphate medium containing 50% of the standard concentration; and 5P represents high-phosphate medium containing fivefold the standard concentration. Data represent mean ± SD from four independent experiments. **D)** Representative culture images. **E)** Growth curves. **F)** Final OD. **G)** Changes in chlorophyll and carotenoid content. **H)** The intracellular reactive oxidative species (ROS) of cells from **D**. ROS levels were normalized to the values of LT under 1P condition. Data shows mean ± SD from four independent experiments as in **D**. In **F -H**, statistical significance (p < 0.05) is indicated by different letters above error bars, as determined by one-way ANOVA with *post-hoc* Tukey HSD test. **ALT TEXT:** Panels A–C show that overexpression of the Pho (phosphate) regulon members, PstS1 and Sll0654, enhances high-light tolerance. Additional phosphate supplementation experiments in panels D–H demonstrate that phosphate availability affects growth, pigment accumulation, and reactive oxygen species levels (ROS) during high-light acclimation.

The up-regulation of Pho regulon, together with the improved growth of both OE lines under moderate HL, suggests that enhanced P_i_ acquisition is important for HL-acclimation. Because elevated light intensity increases photosynthetic energy input and then stimulates a series of highly P_i_-demanding cellular metabolisms, including nucleotides synthesis, ATP turnover, protein repair, membrane remodeling, and generation of phosphorylated intermediates during the CBB cycle (Bhaya et al., 2000; Pitt et al., 2010; Hernandez and Munne-Bosch, 2015), we hypothesized that P_i_-availability may become limiting during high light. To test this possibility, we tested the effects of the growth of LT and two OE lines in media containing different P_i_ regimes. Under Pi-depleted condition, LT and the two OE lines exhibited pronounced growth defects under CL and mHL’, accompanied by strongly reduced pigment accumulation (**Supplementary Fig. S11A, B**). These observations indicate that external phosphate is required for sustained growth and suggest that the improved performance of both OE lines under moderate HL, as well as their growth trade-off under CL, is closely linked to phosphate availability.

We next compared the growth of LT and both OE lines in standard BG11 medium (1P), low P_i_ medium containing 50% of the standard concentration (0.5P), and high Pi medium containing fivefold more than the standard concentration (5P). Under 0.5P condition, LT displayed slower growth and lower pigment accumulation than both OE lines under mHL′, resembling the phenotype observed under standard 1P condition **(Fig. 6D–G)**. In contrast, the growth retardation of LT under mHL′ was largely rescued through P_i_ supplementation under 5P condition, and no obvious differences in growth curves or final OD values were observed between LT and the OE lines (**Fig. 6D–E**). Nevertheless, PstS1-OE still maintained a higher chlorophyll content than LT (**Fig. 6G**). Notably, the growth and pigment contents of both OE lines were relatively stable across the three P_i_ conditions (**Fig. 6D–G**), except for a slight decrease in pigment content in PstS1-OE under 0.5P condition (**Fig. 6G**). Taken together, these results indicate that standard phosphate availability may become insufficient to support the metabolic capacity required for growth under moderate HL, whereas enhanced expression of phosphate-acquisition components improves growth performance even under less P_i_ condition (0.5P), likely by alleviating phosphate-related constraints.

We further determined the intercellular ROS content under different P_i_ conditions as a proxy for HL-induced oxidative stress. Interestingly, the ROS content significantly decreased in LT and the OE lines under the 5P condition (**Fig. 6H**), although no obvious improvement of growth was observed in OE lines with additional P_i_ supplementation. This suggests that adequate phosphate supply may help mitigate ROS accumulation under HL, possibly by supporting metabolic flux and preventing redox imbalance caused by excess accumulation of reducing power. Under the 0.5P condition, ROS levels were also reduced in the OE lines, whereas no comparable decrease was observed in LT (**Fig. 6H**), corresponding to its impaired growth. This observation is consistent with previous findings that phosphate limitation can induce PhoB-dependent ROS detoxification pathways (Qiu et al., 2024). To further investigate the effect of stronger P_i_ limitation, we cultivated LT and both OE lines in medium containing only 20% of the standard phosphate concentration (0.2P). Unexpectedly, LT reached a growth and final OD comparable to that of the OE lines under mHL′, indicating that the apparent growth defect of LT was reduced under 0.2P. However, all three lines became more brownish and accumulated substantially lower pigment levels (**Supplementary Fig. S11C–G**), suggesting that the increased biomass did not reflect fully restored physiological performance but rather a distinct severe-phosphate-limitation acclimation state. Consistently, ROS levels were markedly decreased in all three lines under mHL′ with 0.2P (**Supplementary Fig. S11F**). Collectively, these results indicate that both phosphate supplementation and strong phosphate limitation can reduce ROS accumulation under moderate HL, but likely through different mechanisms.

In addition, we compared the growth of LT and both OE lines under CL across different P_i_ concentrations to determine whether the growth trade-off of OE lines can be mitigated through modulating P_i_ availability. However, both OE lines generally grew more slowly than LT under all tested Pi conditions, with the exception of a slight improvement in Sll0654-OE under 0.5P conditions (**Supplementary Fig. S11H, I**). Chlorophyll contents were largely similar among the three lines, except under 0.2P condition, where its accumulation decreased (**Supplementary Fig. S11J**). By contrast, carotenoid contents remained higher in both OE lines than in LT across all tested P_i_ concentrations (**Supplementary Fig. S11J**). These results indicate that the growth trade-off under CL was not rescued by either reducing or increasing P_i_ availability, suggesting that this phenotype is not simply caused by external phosphate limitation.

## Discussion

The development of HL tolerant strains through ALE has unveiled a complex genetic landscape with various combinations of point mutations (Dann et al., 2021). Achieving tolerance to HL conditions requires the synergistic effect of multiple genetic alterations, with mutations in different genes leading to similar epistatic outcomes. In strain UMMM2, mutations NdhF1_F124L_ and EF-G2_R461C_ co-occur, but NdhF1_F124L_ appears at a low frequency (see **Fig. 1**). When combined, these mutations show no additive effect on growth under various light conditions, with the negative effect of EF-G2_R461C_ on CEF prevailing (Dann et al., 2021). This lack of synergy suggests functional redundancy or slight antagonism in their effects on HL tolerance, explaining why they do not fully segregate together in ALE strains.

Although this study focused on the non-motile LT background in which the HL-tolerant mutations were originally selected, it will be important in future work to test whether the ALE-derived mutations also confer physiological benefits in the motile WT background. Such experiments, however, will require careful interpretation because WT cells can partly avoid HL stress by moving toward shaded regions within the culture vessel. Therefore, analyses in the WT background should distinguish between cell-intrinsic HL acclimation and photoprotection resulting from motility. Moreover, the genomic differences between LT and WT have been analyzed in detail in our recent study (Figueroa-Gonzalez et al., 2026). In total, 37 mutations were identified in LT relative to WT, including 29 chromosomal mutations and 8 plasmid mutations. None of these LT-specific mutations has been directly demonstrated to confer HL-tolerance, and that no overlap was found between these mutations and the HL-tolerance-associated mutations identified in the HL-ALE experiment. Therefore, while these genomic differences may contribute to the distinct behavior of LT compared with WT, we cannot currently assign the difference in HL tolerance to a specific candidate mutation.

Proteomics analysis revealed a gradual increase in the expression of ten NDH-1 subunits from CL to HL conditions in sHL-tolerant strains, peaking at different light intensities for different strains (see **Supplementary Fig. S6E**). The NdhF1_F124L_ mutation showed downregulation of *ndhF1* transcripts under various light conditions (see **Supplementary Fig. S3**), suggesting a shift towards NDH-1MS complexes containing NdhF3, which may facilitate CEF and CO_2_ uptake during extended HL stress (Bernat et al., 2011; Zhang et al., 2020). To fully understand the mechanisms at play, it remains to be determined whether the effects of mutating NdhF1 are specific or if similar results could be achieved by mutating NdhD1 proteins, which are present in NDH-1L but replaced by NdhD3 in NDH-1MS. Moreover, it is crucial to investigate whether the NdhF1 mutation affects the accumulation of NdhF1 itself and induces the shift of different NDH-1 complexes, and how this mutation leads to increased overall accumulation of NDH proteins, potentially resulting in higher CEF.

Cyanobacteria typically downregulate photosynthetic complex subunits under HL to prevent excessive photodamage (Muramatsu and Hihara, 2012; Abdel-Salam et al., 2023), and a recent genome-wide CRISPR interference study revealed that PBS repression, while generally detrimental, is beneficial under HL (Miao et al., 2023). The most notable effect of the EF-G2_R461C_ mutation is the reduction in PBS size, primarily due to CpcC2 linker protein depletion, which mitigates excess light energy absorption (see **Fig. 5**). This adaptation aligns with findings in microalgae, where reducing the light-harvesting antenna cross-section improved growth under HL conditions (Kirst et al., 2014; Lea-Smith et al., 2014; Negi et al., 2020). Although loss of CpcC2 improved high-light acclimation, Δ*cpcC2* did not fully phenocopy the sHL-tolerance of EF-G2_R461C_. This is consistent with the previous study, in which reduced photoinhibition and improved growth under sHL were observed in a mutant lacking both *cpcC2* and *cpcC1*, but not in the *cpcC2* single mutant (Lea-Smith et al., 2014). Therefore, additional regulatory mechanisms are likely required to achieve the pronounced sHL-tolerance as observed in EF-G2_R461C_. The EF-G2_R461C_ mutation induces an HL-like response even under CL conditions, resulting in reduced light energy input compared to other strains when exposed to strong HL. This adaptation, while protective under extreme HL, is not advantageous under lower light intensities due to limited light energy absorption, consistent with previous findings in a *Synechocystis* CpcC1C2-deficient strain (Lea-Smith et al., 2014).

Typically, intense illumination causes oxidative inactivation of elongation factors, suppressing protein synthesis and inhibiting photosystem II repair (Kojima et al., 2007; Kojima et al., 2009). The suggestion that EF-G2_R461C_ mutation confers protection by adding a ROS-sensitive cysteine is controversial, given previous research showing increased photoprotection when replacing cysteine with serine in EF-G1 (Ejima et al., 2012). A plausible explanation could be that the addition of a cysteine in EF-G2_R461C_ might increase its vulnerability to ROS stress. Future studies should focus on examining *de novo* protein synthesis and PSII repair in EF-G2_R461C_ under sHL conditions.

HL conditions induce the expression of various proteins, including high light-inducible polypeptides (Hlips), CO_2_-concentrating mechanism (CCM) proteins, the Pho regulon, and NDH subunits (He et al., 2001; Hihara et al., 2001; Suzuki et al., 2004; Pitt et al., 2010; Battchikova et al., 2011; Muramatsu and Hihara, 2012; Long et al., 2016; Zhang et al., 2020). EF-G2_R461C_ primarily upregulates Hlips, CCM proteins, and CCM proteins, with NdhF1_F124L_ showing a lesser effect (see **Supplementary Fig. S6**). The Pho regulon is induced specifically by EF-G2_R461C_ (see **Supplementary Fig. S8A**). Maximum NDH-1 subunit induction is observed with NdhF1_F124L_ at mHL and EF-G2_R461C_ at sHL (see **Supplementary Fig. S6E**). Despite these varied responses, that individual adaptations are insufficient to confer sHL tolerance. This is evidenced by the sHL intolerance of NdhF1_F124L_ and our experiments overexpressing Pho regulon genes (see **Fig. 6**). The greater potency of EF-G2_R461C_ in conferring sHL tolerance suggests that a combination of multiple adaptive mechanisms is necessary for the observed HL tolerance in our evolved strains. Thus, future work should aim to identify these cooperating mechanisms, for example by testing the cooperation of *cpcC2*-deletion and overexpressing Pho regulon or by assessing whether additional deletion of CpcC1 in Δ*cpcC2* further enhances the phenotype, since CpcC1 also showed reduced accumulation, although to a lesser extent than CpcC2.

HL increases photosynthetic electron flow, ATP turnover and carbon fixation, which likely enhances the demand for P_i,_ one of the most abundant macro nutrients incorporated in to multiple macro biomolecules, and therefore creates a relative P_i_ limitation and trigger a phosphate-starvation-like response (Pitt et al., 2010; Hernandez and Munne-Bosch, 2015). Consistently, improved growth of both OE lines was observed under 50% and standard P_i_ conditions, but this advantage disappeared under fivefold P_i_ supplementation (see **Fig. 6** and **Supplementary Fig. S11**). These results suggest that phosphate availability becomes limiting during moderate HL acclimation and that enhanced expression of P_i_ acquisition components can be beneficial when P_i_ is available but potentially insufficient to meet the increased metabolic demand. At 20% P_i_ LT reached a high apparent OD under moderate HL, but the brownish phenotype and reduced pigment accumulation indicate HL-compensatory physiological impairment rather than true healthy acclimation. This suggests that severe phosphate limitation may suppress ROS accumulation by reducing photosynthetic electron flow and activating antioxidant defenses, whereas sufficient phosphate may support metabolic activity and redox balance. By contrast, the growth trade-off of both OE lines under CL was not rescued by altering external P_i_ availability (see **Supplementary Fig. 11**), suggesting that perturbation of phosphate homeostasis caused by constitutive activation of P_i_-acquisition components may create a metabolic or regulatory burden under CL, where phosphate demand is relatively low.

In sum, EF-G2 appears to be a potential hub for HL adaptation, being mutated in all HL-ALE strains (Dann et al., 2021). The mechanisms by which NdhF1 and EF-G2 mutations affect NDH complexes and CpcC2 accumulation remain unclear. Although our results demonstrate the impact of NdhF1_F124L_ on NDH-1 subunit abundance, direct biochemical analysis, for example by 2D gel electrophoresis, is still required to examine the accumulation and composition of various NDH-1 complexes, thereby further elucidating the mechanism underlying the HL-tolerance of the NdhF1 mutation. Given the role of EF-G in translation elongation, future experiments such as polysome profiling or ribosome profiling will be required to determine the effect of EF-G2_R461C_ on global or transcript-specific translation. HL-ALE produced strains with mutations in key proteins resulting in maximal tolerance, and also endowed the cells with novel regulatory approaches to employ the universal HL stress alleviating mechanisms. This adaptation strategy allows identification of unknown hubs for rewiring transcriptional and translational networks, facilitating the development of strategies to enhance photosynthetic efficiency.

## Author contributions

D.L. and W.C. conceived the project. D.L. provided funding. D.L. and W.C. designed all experiments. W.C. performed the experiments with support of M.D., C.O. and S.S. in the proteomics experiment. E.A.S. helped with the transcriptomics analysis and interpretation of the results. D.L. and W.C. wrote the manuscript. All authors read and approved the final manuscript.

## Acknowledgements

Open Access funding enabled and organized by Projekt DEAL. The artificial intelligence tool ChatGPT was used to improve readability and language.

## Supplementary information

The online version contains supplementary material available at xxxx.

## Data availability

RNA-Seq data were deposited in the Gene Expression Omnibus repository with the accession number GSE268898. Proteomics data have been deposited with the ProteomeXchange consortium via the PRIDE (Perez-Riverol et al., 2022) partner repository under accession number PXD052516. All additional data are available upon request from the corresponding author.

