## Supplemental Tables for "Multi-omics dissection of transcriptional and post-transcriptional responses in cyanobacterial high-light adaptation"

**Supplementary Table S1. Common HL-tolerance related genes (HL-DEGs) in the HL-tolerant strains.**

| **Gene ID** | | **Old locus tag** | |  | **Name** | **Description** |
| --- | --- | --- | --- | --- | --- | --- |
| **Down_regulated HL-DEGs** | | |  |  |  |  |
|  | SGL_RS16410 | | sll0743 |  | sll0743 | hypothetical protein |
|  | SGL_RS16215 | | slr0699 |  | slr0699 | unknown protein |
|  | SGL_RS08830 | | sll1596 |  | kaiB2 | circadian clock protein KaiB homolog |
|  | SGL_RS08295 | | slr0870 |  | slr0870 | hypothetical protein |
|  | SGL_RS05160 | | slr1396 |  | slr1396 | unknown protein |
|  | SGL_RS01830 | | slr6101 |  | slr6101 | plasmid-encoded putative toxin/antitoxin pair |
|  | SGL_RS01700 | | slr6072 |  | slr6072 | unknown protein |
|  | SGL_RS01575 | | slr6045 |  | slr6045 | unknown protein |
|  | SGL_RS01430 | | slr6013 |  | slr6013 | unknown protein |
|  | SGL_RS01030 | | sll7086 |  | sll7086 | unknown protein |
|  | SGL_RS01025 | | sll7085 |  | sll7085 | unknown protein |
|  | SGL_RS00605 | | sll5128 |  | sll5128 | unknown protein |
|  | SGL_RS09525 | |  |  |  | hypothetical protein |
|  | SGL_RS14650 | |  |  |  | type II toxin-antitoxin system HicA family |
| **Up_regulated HL-DEGs** | | |  |  |  |  |
|  | SGL_RS16520 | | sll0037 |  | cbiX | sirohydrochlorin cobaltochelatase |
|  | SGL_RS12205 | | sll0141 |  | sll0141 | putative periplasmic adaptor protein |
|  | SGL_RS13285 | | sll0314 |  | sll0314 | periplasmic protein, function unknown |
|  | SGL_RS14445 | | sll0381 |  | sll0381 | hypothetical protein |
|  | SGL_RS14430 | | sll0384 |  | sll0384 | cation and iron carrying protein |
|  | SGL_RS14425 | | sll0385 |  | sll0385 | ATP-binding protein of ABC transporter |
|  | SGL_RS15490 | | sll0477 |  | sll0477 | putative biopolymer transport ExbB-like protein |
|  | SGL_RS14310 | | sll0608 |  | ycf49 | hypothetical protein YCF49 |
|  | SGL_RS03825 | | sll0656 |  | nucH | extracellular nuclease |
|  | SGL_RS17785 | | sll0720 |  | sll0720 | RTX toxin activating protein homolog |
|  | SGL_RS17775 | | sll0722 |  | sll0722 | [unknown protein](https://www.sciencedirect.com/topics/chemistry/sirohydrochlorin) |
|  | SGL_RS17765 | | sll0723 |  | sll0723 | unknown protein |
|  | SGL_RS07350 | | sll1308 |  | sll1308 | probable oxidoreductase |
|  | SGL_RS17545 | | sll1483 |  | sll1483 | periplasmic protein, similar to   transforming growth factor induced protein |
|  | SGL_RS04105 | | sll1507 |  | sll1507 | salt-induced periplasmic protein |
|  | SGL_RS17790 | | sll1552 |  | sll1552 | unknown protein |
|  | SGL_RS06130 | | sll1968 |  | pmgA | photomixotrophic growth related protein |
|  | SGL_RS00225 | | sll5050 |  | sll5050 | probable glycosyltransferase |
|  | SGL_RS00580 | | sll5123 |  | sll5123 | SOS mutagenesis and repair, UmuD protein homolog |
|  | SGL_RS01415 | | sll6010 |  | sll6010 | unknown protein |
|  | SGL_RS01685 | | sll6069 |  | sll6069 | unknown protein |
|  | SGL_RS14105 | | slr0492 |  | menE | O-succinylbenzoic acid-CoA ligase |
|  | SGL_RS15495 | | slr0513 |  | futA2 | iron transport system substrate-binding protein |
|  | SGL_RS15505 | | slr0516 |  | slr0516 | hypothetical protein |
|  | SGL_RS02010 | | slr1485 |  | slr1485 | putative phosphatidylinositol phosphate kinase |
|  | SGL_RS09605 | | slr2131 |  | acrB | RND multidrug efflux transporter |
|  | SGL_RS00245 | | slr5054 |  | slr5054 | probable glycosyltransferase |
|  | SGL_RS01405 | | slr6008 |  | slr6008 | unknown protein |
|  | SGL_RS00880 | | ssl7053 |  | ssl7053 | hypothetical protein |
|  | SGL_RS04605 | | ssr3465 |  | ssr3465 | unknown protein |
|  | SGL_RS19830 | |  |  |  | hypothetical protein |

Commonly identified up- and down-regulated HL-DEGs in HL-tolerant WT, UMMM2, and EF-G2_R461C_/ NdhF1_F124L_+EF-G2_R461C_ as showed in **Fig. 2C** are listed.

**Supplementary Table S2. Common HL-tolerance related proteins (HL-DEPs) in the HL-tolerant strains.**

|  | **Protein ID** | **Name** | | **Uniprot ID** | **Description** |
| --- | --- | --- | --- | --- | --- |
| **Down_regulated HL-DEPs** | | |  |  |  |
|  | sll0172 | | sll0172 | Q55558 | periplasmic protein, function unknown |
|  | sll0258 | | psbV | Q55013 | cytochrome c550 |
|  | sll0293 | sll0293 | | Q55547 | unknown protein |
|  | sll0915 | | pqqE | P74305 | periplasmic protease |
|  | sll1491 | | sll1491 | P74598 | periplasmic WD-repeat protein |
|  | sll1835 | | sll1835 | P73111 | periplasmic protein, function unknown |
|  | slr1266 | | slr1266 | P74179 | hypothetical protein |
|  | slr1274 | | pilM | P74186 | probable fimbrial assembly protein PilM |
|  | slr1276 | | slr1276 | P74188 | hypothetical protein |
| **Up_regulated HL-DEGs** | | |  |  |  |
|  | sll1394 | | msrA1 | P72622 | peptide methionine sulfoxide reductase |
|  | slr1367 | | glgP | P73546 | glycogen phosphorylase |
|  | sll1961 | | sll1961 | P73804 | hypothetical protein |
|  | sll0553 | | sll0553 | Q55390 | hypothetical protein |
|  | slr0453 | | slr0453 | P74690 | hypothetical protein |
|  | slr2075 | | groES | Q05971 | 10kD chaperonin |
|  | sll1294 | | pilJ | P73173 | methyl-accepting chemotaxis protein |
|  | slr0607 | | slr0607 | P74754 | hypothetical protein |
|  | ssl1633 | | hliC, scpB | P73563 | high light-inducible polypeptide HliC |
|  | slr0884 | | gap1 | P49433 | glyceraldehyde 3-phosphate dehydrogenase 1 |
|  | sll0680 | | pstS1 | Q55199 | periplasmic phosphate-binding protein |
|  | sll1594 | | ndhR | P73862 | ndhF3 operon transcriptional regulator |
|  | sll0934 | | ccmA | P72864 | carboxysome formation protein CcmA |
|  | sll0329 | | gnd | P52208 | 6-phosphogluconate dehydrogenase |
|  | sll0654 | | sll0654 | P72939 | alkaline phosphatase |
|  | slr0516 | | slr0516 | Q55837 | hypothetical protein |
|  | slr1351 | | murF | P45450 | UDP-N-acetylmuramoylalanyl-D-glutamyl-2 6-  diaminopimelate--D-alanyl-D-alanine ligase |
|  | slr1512 | | sbtA | P73953 | sodium-dependent bicarbonate transporter |
|  | slr1204 | | htrA | P73354 | protease |
|  | slr2073 | | ycf50 | P73376 | hypothetical protein YCF50 |
|  | slr1280 | | ndhK | P19050 | NADH dehydrogenase subunit NdhK |
|  | sll1680 | | sll1680 | P72779 | hypothetical protein |
|  | sll1566 | | ggpS | P74258 | glucosylglycerolphosphate synthase |
|  | sll0185 | | sll0185 | Q55770 | hypothetical protein |
|  | sll0381 | | sll0381 | Q55744 | hypothetical protein |

Commonly identified up- and down-regulated HL-DEPs in HL-tolerant WT, and EF-G2_R461C_/ NdhF1_F124L_+EF-G2_R461C_ as showed in **Fig. 3C** are listed.
