## Supplemental Figures for "Multi-omics dissection of transcriptional and post-transcriptional responses in cyanobacterial high-light adaptation"

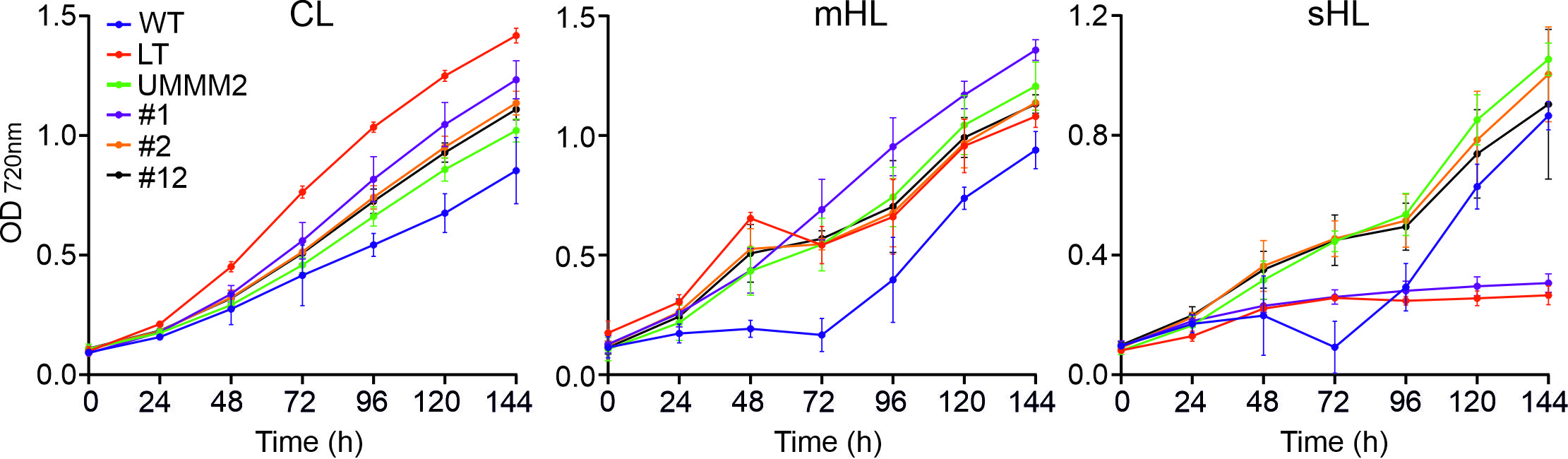

**Supplementary Fig. S1.** **Growth curves of WT, LT, UMMM2, NdhF1_F124L_ (#1), EF-G2_R461C_ (#2), and NdhF1_F124L_+EF-G2_R461C_ (#12) grown for a week under varying light intensities (CL, mHL, and sHL)**. Optical density (OD_720 nm_) was measured hourly using the built-in real-time monitor system in Multi-Cultivator OD-1000 devices. For clarity, values at 24-hour intervals were plotted. Data shows mean ± SD from three independent experiments as in **Fig. 1**.

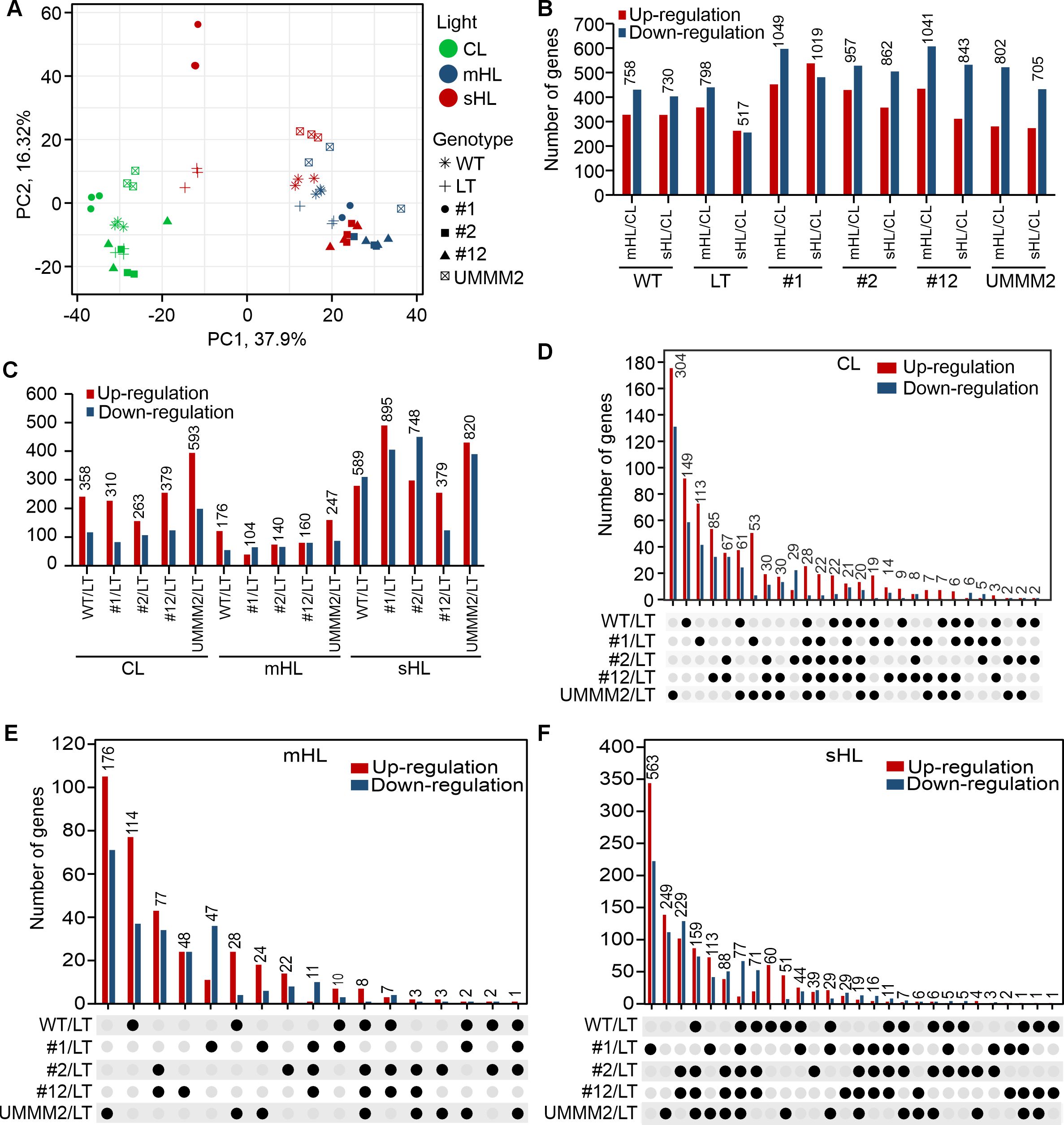

**Supplementary Fig. S2. Detailed transcriptomic analyses.** **A)** Principal Component Analysis of CPM-normalized read counts for all identified transcripts across various genotypes and light conditions. The plot displays the two principal components with the highest variance percentages. One NdhF1_F124L_ (#1) sample at mHL (blue circle) was excluded due to its significant deviation from the other two replicates. Two NdhF1_F124L_ (#1) samples at sHL (red circle) were highly similar and appear as a single point on the plot. **B)** This panel shows the number of Differentially Expressed Genes (DEGs) for mHL vs. CL and sHL vs. CL in WT, LT and the four mutant strains: NdhF1_F124L_ (#1), EF-G2_R461C_ (#2), NdhF1_F124_+EF-G2_R461C_ (#12) and UMMM2. **C)** The number of DEGs for WT and the four mutant strains vs. LT in CL, mHL and sHL conditions is presented. **D-F)** UpSet plot illustrating the number of commonly up- or down-regulated DEGs in various intersections of WT and four mutant strains compared to LT under CL **(D)**, mHL **(E)**, and sHL **(F)** conditions. The matrix below the x-axis shows intersections between the five genotypes relative to LT, with black dots indicating which genotypes are part of an intersection. The y-axis shows the number of up- or down-regulated transcripts in each intersection. Up- and down-regulated genes are analyzed separately but combined in the plot for easier visualization. Intersections are ordered by the total number of regulated genes (shown at the top of each set).

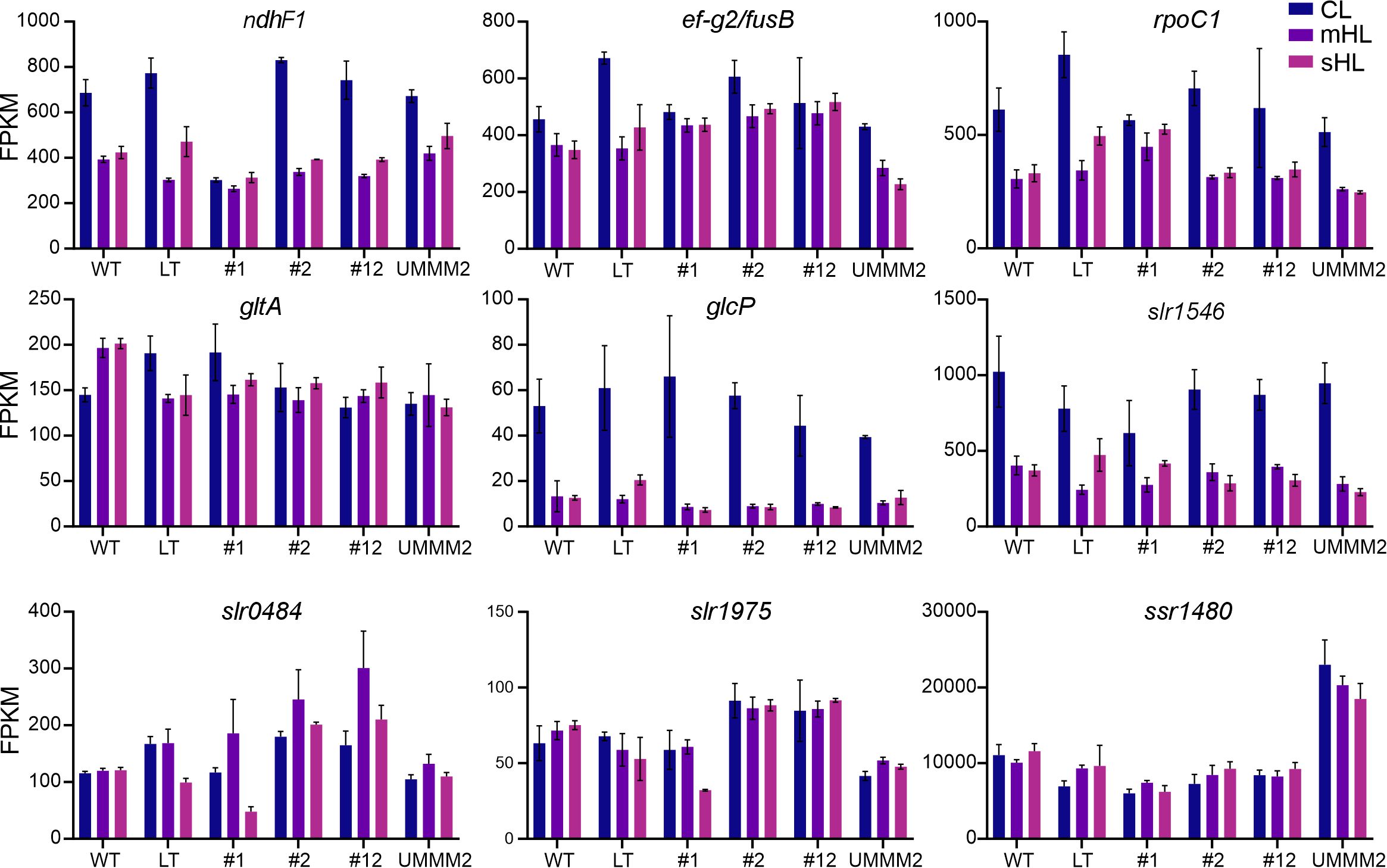

**Supplementary Fig. S3. Transcript abundance of genes mutated in UMMM2 in different genotypes and light conditions.** This figure displays transcript abundance (FPKM) for nine genes with high-frequency mutations in UMMM2, shown for WT, LT, NdhF1_F124L_ (#1), EF-G2_R461C_ (#2), NdhF1_F124L_+EF-G2_R461C_ (#12) and UMMM2 cells under three different light intensities.

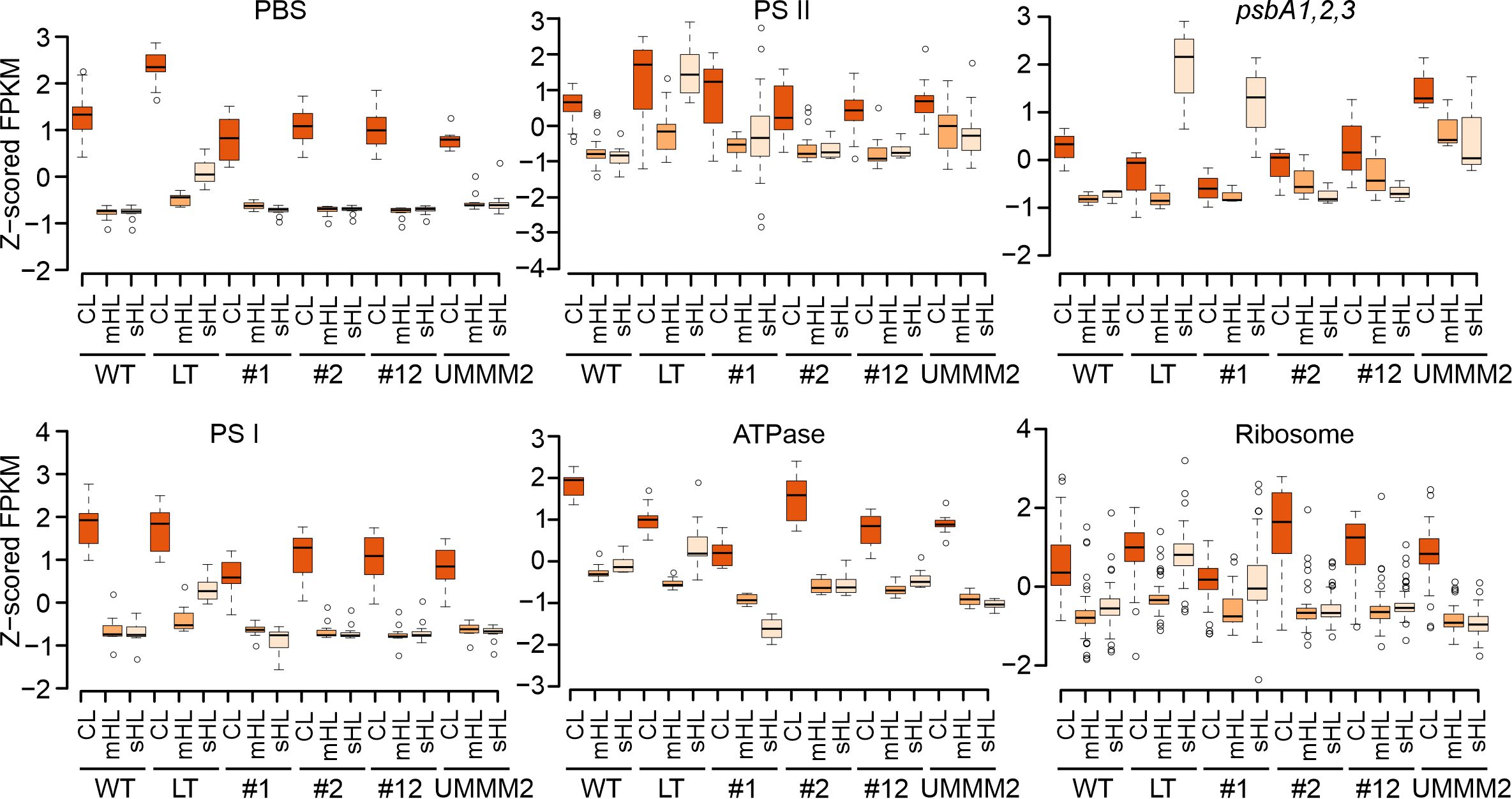

**Supplementary Fig. S4. Overview of transcriptional responses of photosynthesis- and ribosome-related genes in different genotypes and light conditions.** Z-scored FPKM values are presented for WT, LT, NdhF1_F124L_ (#1), EF-G2_R461C_ (#2), NdhF1_F124L_+EF-G2_R461C_ (#12) and UMMM2 cells under three different light intensities. The data covers five gene sets (all phycobilisome, PSII, PSI, ATP synthase, and ribosome subunits) and the three *psbA* genes.

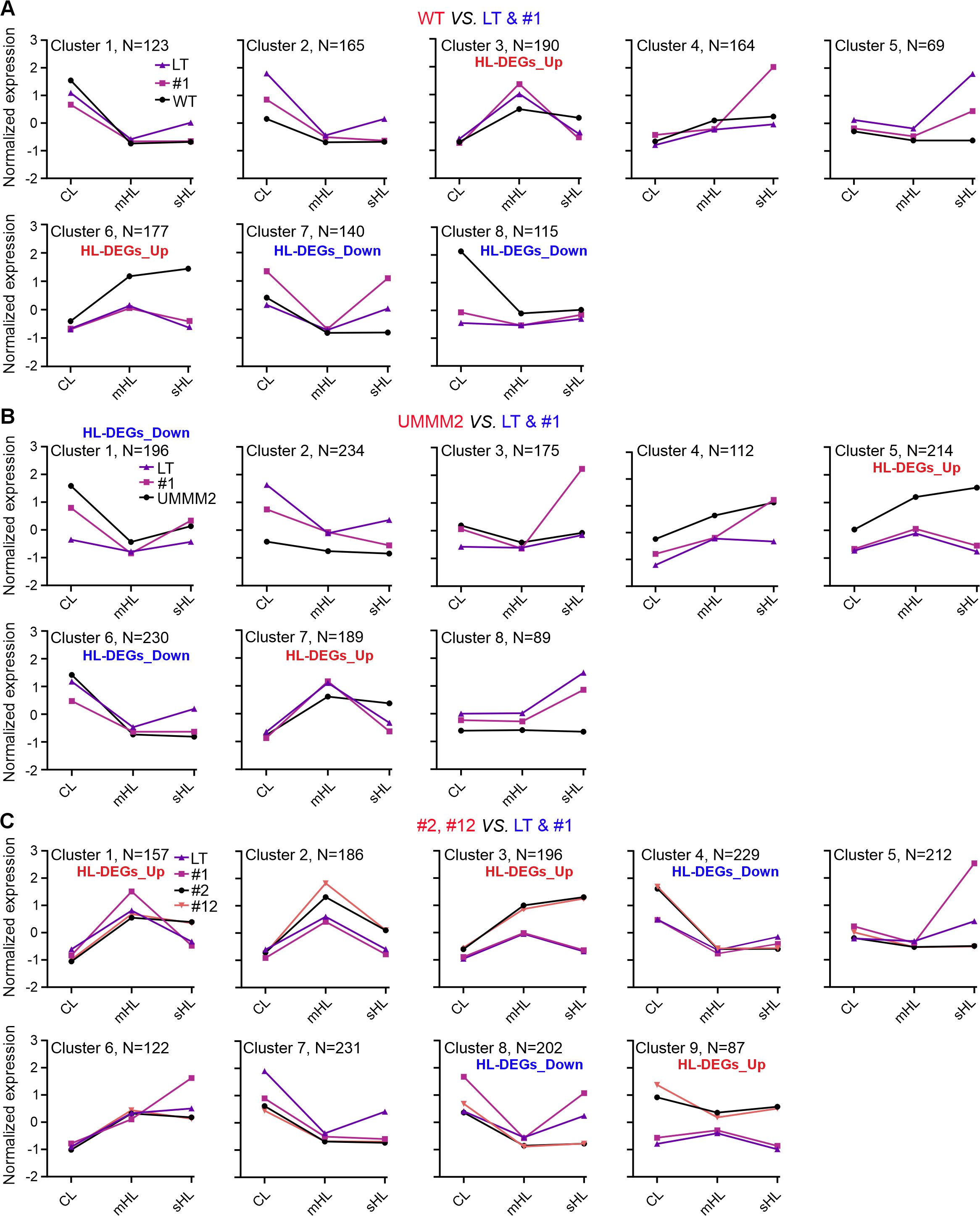

**Supplementary Fig. S5.** **Comparison of DEG clusters between HL-tolerant and -intolerant strains.**

**A-C)** Line plots show averaged gene expression patterns from Fuzzy c-means soft clustering analysis, comparing sHL-tolerant strains (WT, UMMM2, EF-G2_R461C_ (#2), NdhF1_F124L_+EF-G2_R461C_ (#12)) with intolerant strains (LT, NdhF1_F124L_ (#1)). Comparisons are shown for WT (**A**), UMMM2 (**B**), and #2 and #12 (**C**) against LT and #1. Gene numbers for each group are included. The x-axis represents light intensities (CL, mHL, sHL), and the y-axis shows mean normalized expression.

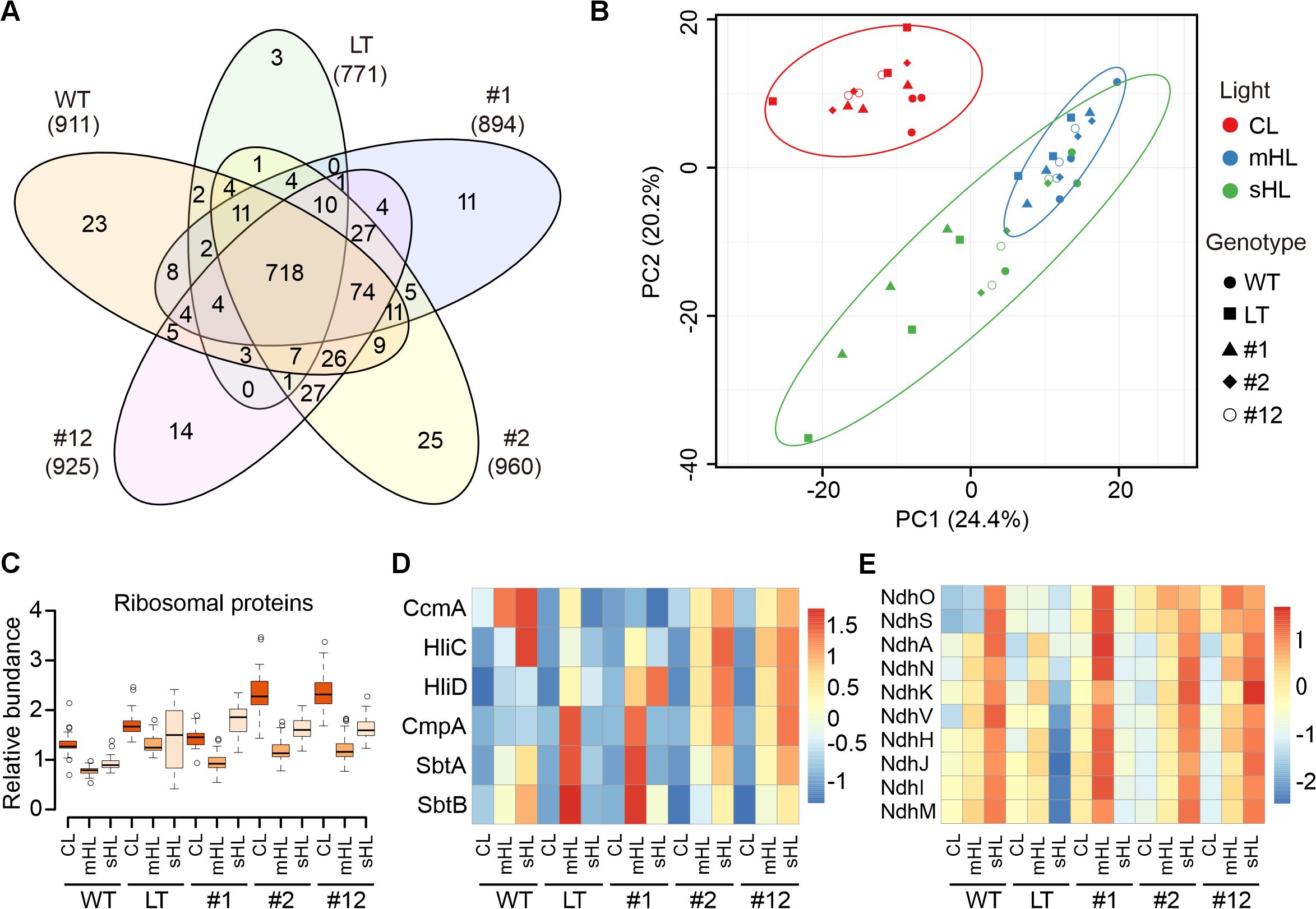

**Supplementary Fig. S6. Overview of proteome changes and specific protein set responses in different genotypes and light conditions.** **A)** Venn diagram showing the overlap and unique proteins accumulated in five strains: WT, LT, NdhF1_F124L_ (#1), EF-G2_R461C_ (#2), and NdhF1_F124L_+EF-G2_R461C_ (#12). Total proteins identified per strain are indicated in parentheses. **B)** Principal Component Analysis of the relative abundance of 718 proteins common to all strains identified in **A**, across different genotypes and light conditions. The plot displays the two principal components with the highest variance percentages. **C-E)** Relative abundance values for WT, LT, UMMM2, NdhF1_F124L_ (#1), EF-G2_R461C_ (#2), and NdhF1_F124L_+EF-G2_R461C_ (#12) cells under three light intensities are shown for three protein sets: ribosome (Rpl1, 5-6, 9-13, 16-21, 23-24, 27, and Rps1, 5-7, 9-10, 12-14, 16, 19-20), Hlip/CCM, and NDH-1 proteins from cluster 12 in **Fig. 3**.

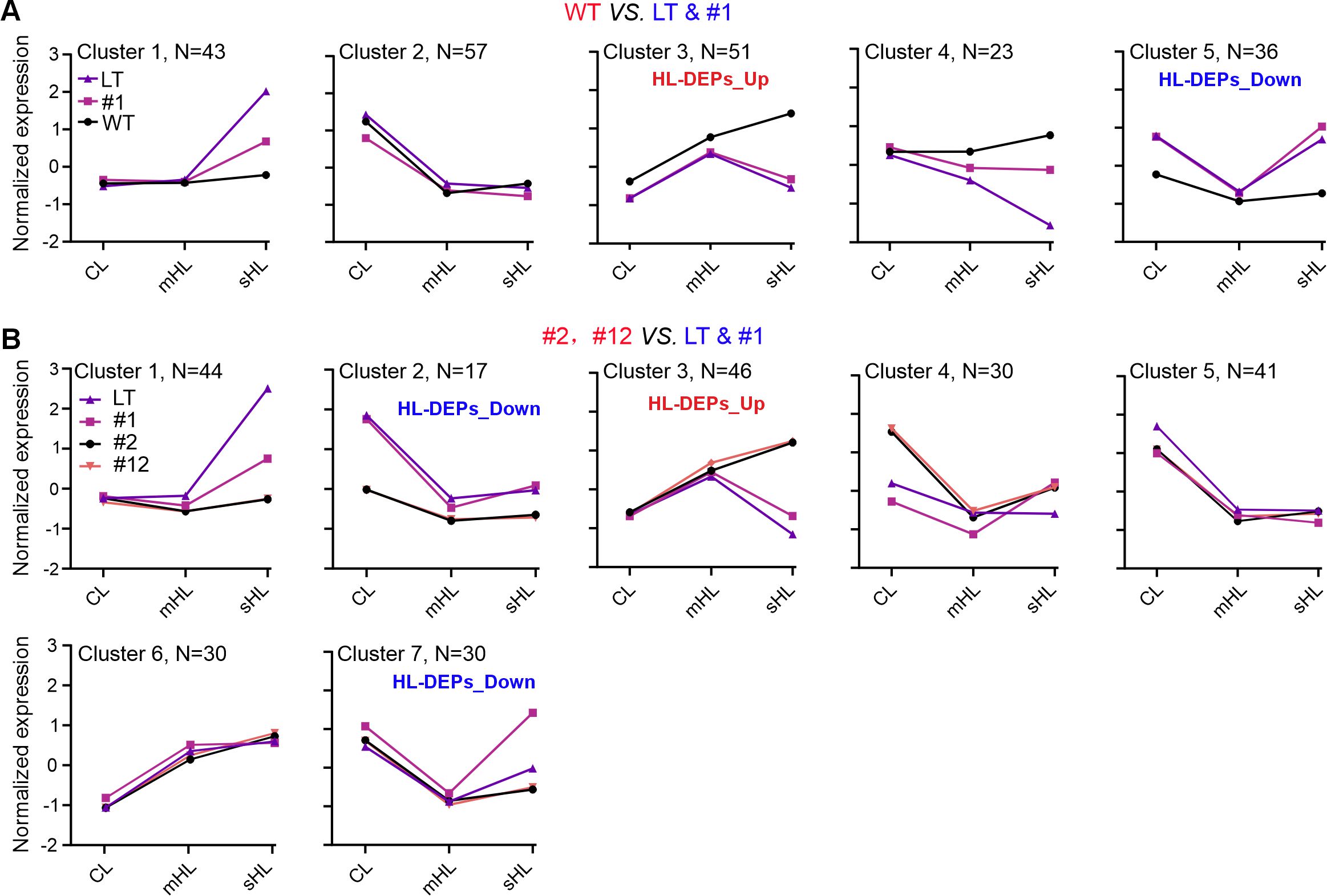

**Supplementary Fig. S7. Comparison of DEP clusters between HL-tolerant and -intolerant strains.**

**A** and **B)** Line plots display averaged protein expression patterns from Fuzzy c-means soft clustering analysis, comparing sHL-tolerant strains (WT, EF-G2_R461C_ (#2), NdhF1_F124L_+EF-G2_R461C_ (#12)) with intolerant strains (LT, Ndh_F1F124L_ (#1)). Comparisons for WT (**A**) and #2 and #12 (**B**) against LT and #1 are shown. Protein numbers for each group are included. The x-axis represents light intensities (CL, mHL, sHL), and the y-axis shows mean normalized expression.

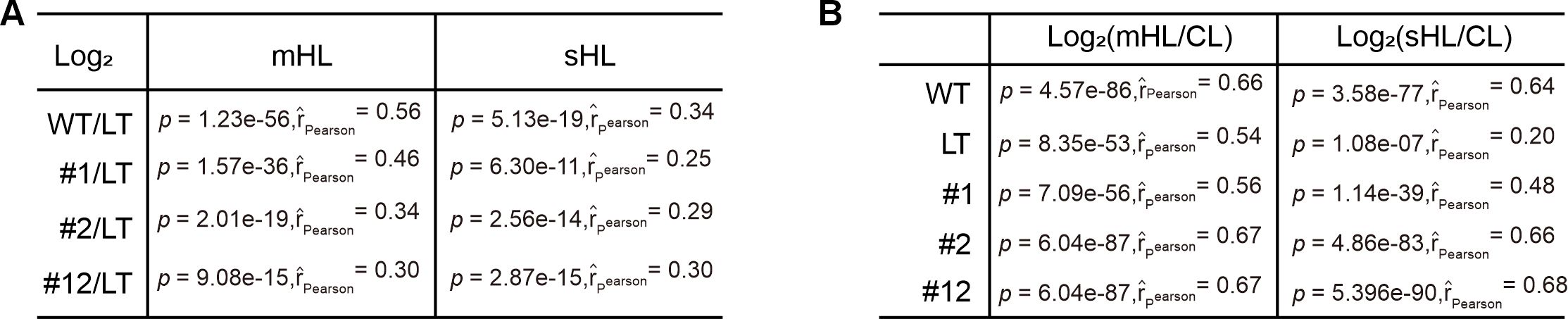

**Supplementary Fig. S8. Comparison of transcriptome and proteome responses.** **A, B)** Correlation analysis between transcriptomes and proteomes. Pearson correlation coefficients (r) and Student t-test p-values (p) are provided for expression changes between all transcript-protein pairs of WT, NdhF1_F124L_ (#1), EF-G2_R461C_ (#2), and NdhF1_F124L_+EF-G2_R461C_ (#12) relative to LT under mHL and sHL conditions (**A**), and between all transcript-protein pairs of mHL and sHL relative to CL for the five genotypes (**B**).

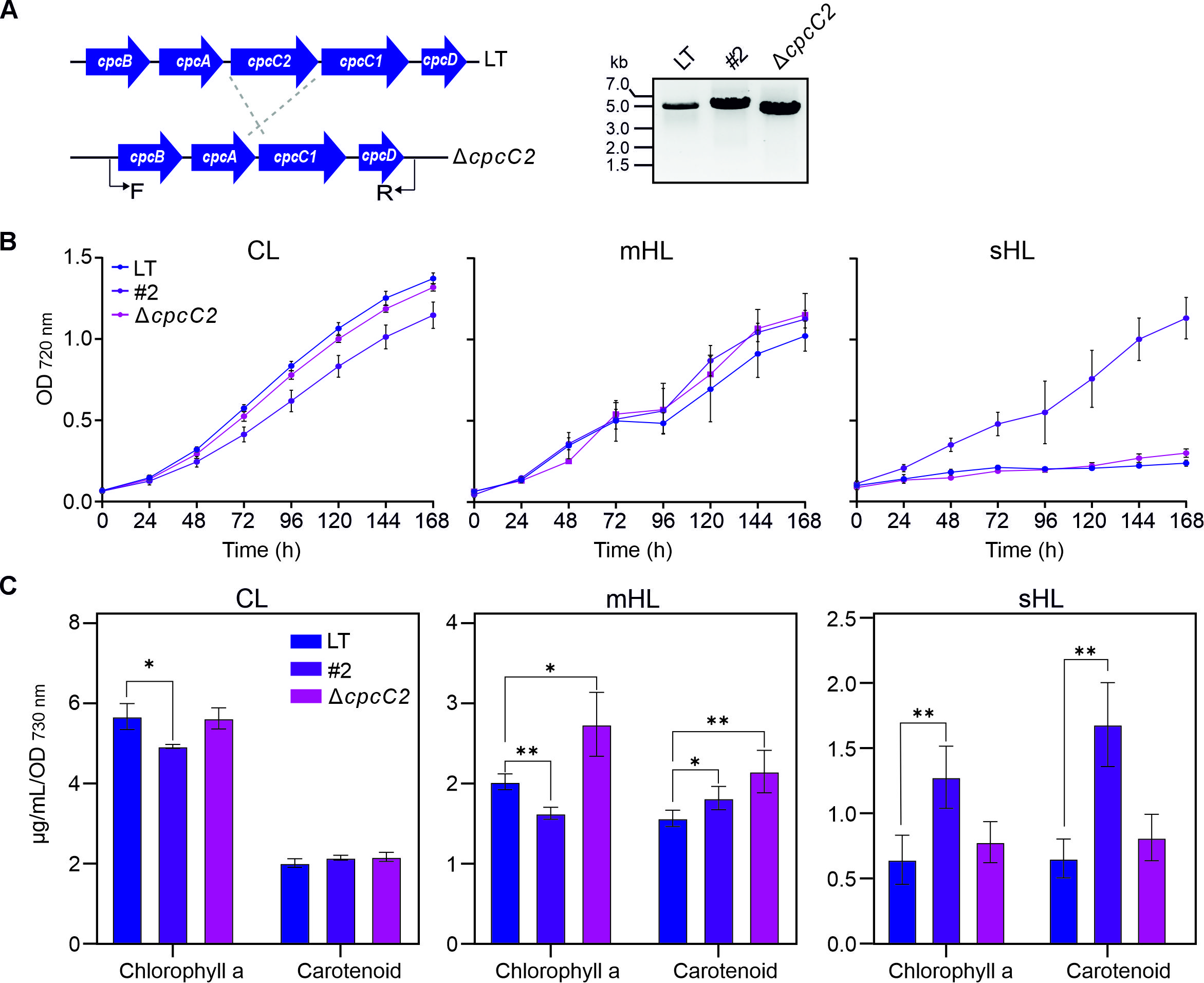

**Supplementary Fig. S9.** **Generation of Δ*cpcC2* and growth analysis. A**) Schematic representation of the construction of the *cpcC2* deletion mutant, Δ*cpcC2*, in the *Synechocystis* LT background. The positions of the forward (F) and reverse (R) primers used for segregation analysis are indicated. Complete segregation was confirmed by genotyping PCR. **B)** Growth curves of LT, EF-G2_R461C_ (#2), and Δ*cpcC2* under CL, mHL, and sHL conditions. Data shows mean ± SD from three independent experiments as in **Fig. 5E**. **C)** Changes in chlorophyll and carotenoid content. Data represents mean ± SD from three independent experiments as in **Fig. 5E**. Statistical significance was determined using two-tailed Student's t-test. *: p < 0.05, **: p < 0.01.

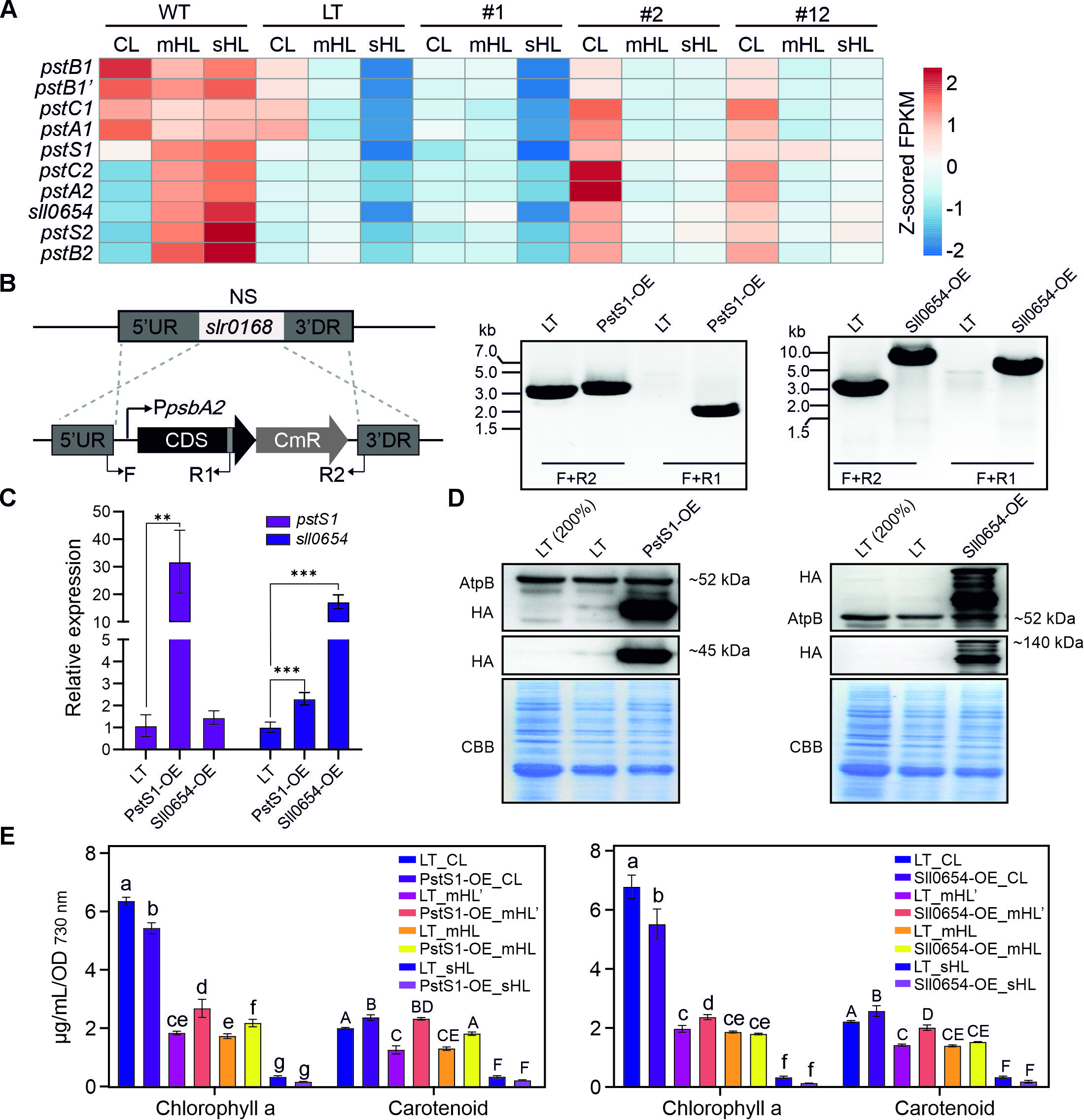

**Supplementary Fig. S10. Generation of overexpression strains and growth analysis. A**) Heatmap of Z-score normalized FPKM for Pho regulon transcripts, comparing sHL-tolerant strains (WT, EF-G2_R461C_ (#2), NdhF1_F124L_+EF-G2_R461C_ (#12)) with sHL-intolerant strains (LT, NdhF1_F124L_ (#1)) under three light conditions. **B**) Outline of the strategy for gene overexpression (OE) in *Synechocystis*. For OE, the target gene, controlled by the *psbA2* promoter (P*_psbA2_*) and linked to a chloramphenicol resistance gene cassette (*CmR*), was inserted into a neutral site (*slr0168*) of the *Synechocystis* LT genome. The diagram indicates the positions of the forward (F1) and reverse (R1, R2) primers used for segregation control. PCR analysis confirmed complete segregation of the OE strains. PstS1-OE denotes the OE of *pstS1*, and Sll0654-OE indicates the OE of *sll0654*. **C**) qRT-PCR analysis of overexpression of *pstS1* and *sll0654* in OE strains. The relative expression was normalized based on the 2^-ΔΔCt^ method. The *rnpB* gene was used as an internal control. Data represents mean ± SD from four independent experiments. Statistical significance was determined using two-tailed Student's t-test. **: p < 0.01, ***: p < 0.001. **D**) Immunoblot analysis of accumulation of HA-fused PstS1 and Sll0654. Approximately 5 µg protein contents were subjected to SDS-PAGE. A Coomassie Brilliant Blue (CBB) stain of the PVDF membrane served as a loading control. **E**) Changes in chlorophyll and carotenoid content. Data represents mean ± SD from three independent experiments as in **Fig. 6A**. Different letters above error bars indicate statistically significant differences (p < 0.05) as determined by one-way ANOVA with post-hoc Tukey HSD test.

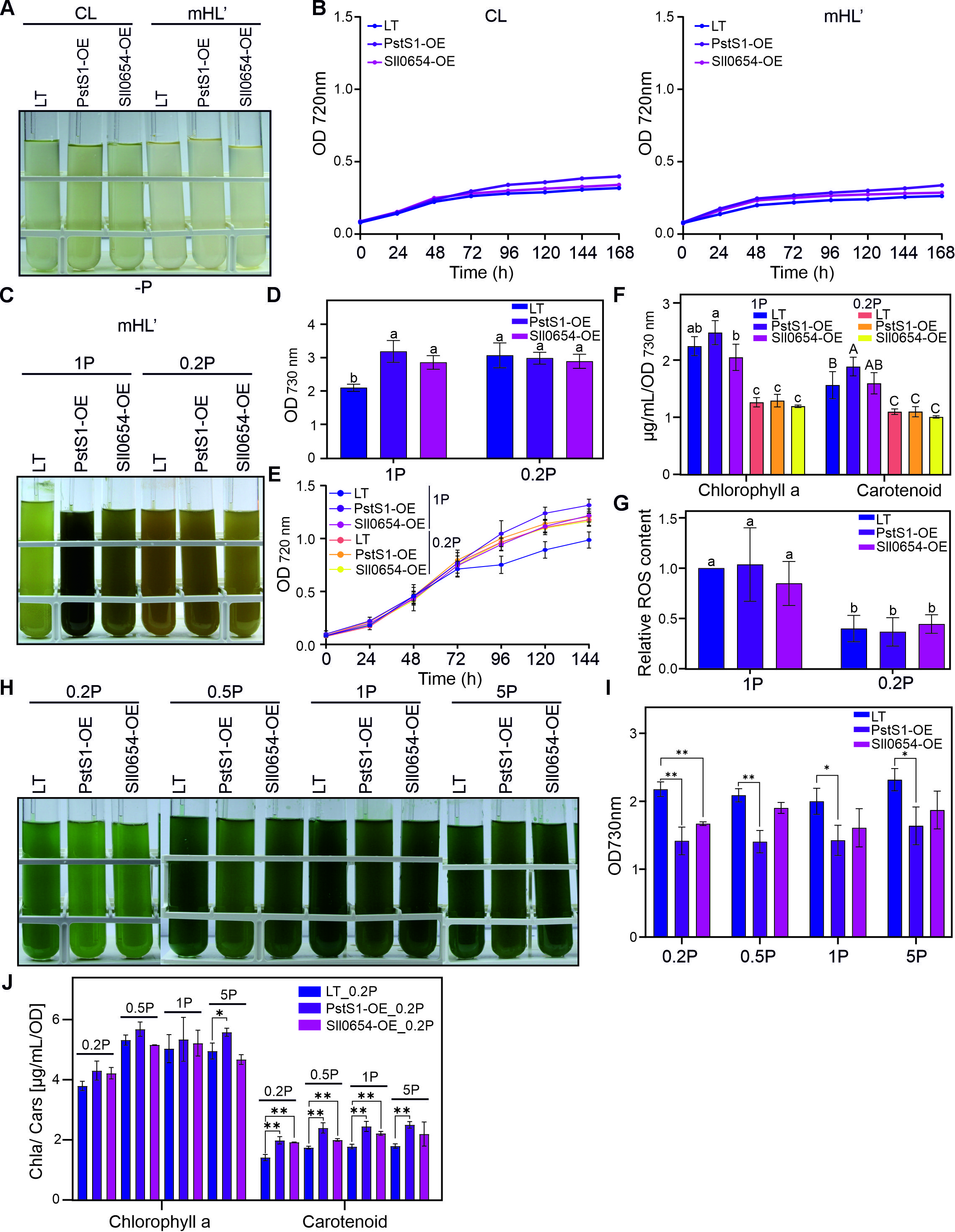

**Supplementary Fig. S11.** **Growth summary of LT, PstS1-OE, and Sll0654-OE different phosphate (P_i_) concentration.** **A**) Culture images of LT, PstS1-OE, and Sll0654-OE grown for 7 days in P_i_-deplete medium under CL and mHL'. **B**) Growth curves of LT, PstS1-OE, and Sll0654-OE grown as in **A**. **C-G)** Growth summary of LT, PstS1-OE, and Sll0654-OE grown for 6 days under mHL' with standard or low P_i_ intensity. 1P represents standard BG11 medium containing 175 μM K₂HPO₄; 0.2P represents low-phosphate medium containing 20% of the standard concentration. Data represent mean ± SD from four independent experiments. **C)** Representative culture images. **D)** Final OD. **E)** Growth curves. **F)** Changes in chlorophyll and carotenoid content. **G)** The intracellular ROS of cells from **(C)**. ROS levels were normalized to the values of LT under 1P condition. Data shows mean ± SD from four independent experiments as in **C**. Statistical significance (p < 0.05) is indicated by different letters above error bars, as determined by one-way ANOVA with post-hoc Tukey HSD test. **H-J)** Growth summary of LT, PstS1-OE, and Sll0654-OE grown for 6 days under CL with low, normal and high P_i_ concentrations. Data represent mean ± SD from three independent experiments, except for Sll0654-OE, for which two independent experiments were performed. **I**) Final OD. **J**) Variation in chlorophyll and carotenoid content. Statistical significance was determined using two-tailed Student's t-test. *: p < 0.05, **: p < 0.01.
